# Single-cell signaling analysis reveals that Major Vault Protein facilitates RasG12C inhibitor resistance

**DOI:** 10.1101/2023.10.02.560617

**Authors:** Jason Z. Zhang, Shao-En Ong, David Baker, Dustin J. Maly

## Abstract

Recently developed covalent inhibitors for RasG12C provide the first pharmacological tools to target mutant Ras-driven cancers. However, the rapid development of resistance to current clinical Ras G12C inhibitors is common. Presumably, a subpopulation of RasG12C-expressing cells adapt their signaling to evade these inhibitors and the mechanisms for this phenomenon are unclear due to the lack of tools that can measure signaling with single-cell resolution. Here, we utilized recently developed Ras sensors to profile the environment of active Ras and to measure the activity of endogenous Ras in order to pair structure (Ras signalosome) to function (Ras activity), respectively, at a single-cell level. With this approach, we identified a subpopulation of KRasG12C cells treated with RasG12C-GDP inhibitors underwent oncogenic signaling and metabolic changes driven by WT Ras at the golgi and mutant Ras at the mitochondria, respectively. Our Ras sensors identified Major Vault Protein (MVP) as a mediator of Ras activation at both compartments by scaffolding Ras signaling pathway components and metabolite channels. We found that recently developed RasG12C-GTP inhibitors also led to MVP-mediated WT Ras signaling at the golgi, demonstrating that this a general mechanism RasG12C inhibitor resistance. Overall, single-cell analysis of structure-function relationships enabled the discovery of a RasG12C inhibitor-resistant subpopulation driven by MVP, providing insight into the complex and heterogenous rewiring occurring during drug resistance in cancer.

## Introduction

The ubiquitous signaling enzyme Ras (<u>Ra</u>t <u>s</u>arcoma virus) is one of the most important oncogenes, with mutations that over activate Ras (GTP-bound) observed in a third of cancers^1^. Due to its strong link to cancer, there have been decades of effort to create therapeutics that target and inhibit mutant Ras. Only recently, have small molecules that specifically target and inhibit mutant Ras been developed. Covalent inhibitors against inactive RasG12C (GDP-bound) (e.g. Sotorasib/AMG-510)^2,3^ have demonstrated promising efficacy and were approved by the FDA in 2022. However, most patients acquire drug resistance^4^. Recently reported RasG12C-GTP inhibitors (RasONi, RMC-6291) are another set of pharmacological tools that seem to more potently inhibit RasG12C tumors^5^, however reactivation of oncogenic signaling is also seen with this inhibitor. This rebound in oncogenic signaling for both drugs begs the question of how resistance against RasG12C inhibitors is achieved, which should presumably be driven by a subpopulation of RasG12C-driven cancer cells. Moreover, it is unclear at the signaling level what differentiates the drug resistant cancer cell population over the responding cells. Many reports identified gene mutations and amplification of components in the MAPK cascade to reactivate the Ras/MAPK pathway^6–9^. Using single-cell RNA sequencing, Xue et al. demonstrated that a subpopulation of cells adapt to RasG12C-GDP inhibition by overexpressing mutant KRas^6^. While single-cell genomic and transcriptomic studies have been performed to understand what differentiates this adaptive cell subpopulation, these changes in DNA and RNA are many steps downstream of Ras inhibition. Therefore, we aimed to perform single-cell analysis on Ras signaling, the most immediate consequence of RasG12C inhibitors, to detect this subpopulation and understand the signaling mechanisms that enable this subpopulation to exist.

Most of these RasG12C inhibitors (both GDP and GTP selective drugs) have been applied to KRasG12C-harboring cancer cells as KRasG12C is seen in various cancer types. While these cells are addicted to KRas and often overexpress mutant KRas after drug application, there is also wildtype (WT) HRas and NRas (H/NRas) inside these cancer cells which can compensate for mutant KRas inhibition and are activated during RasG12C-GDP inhibitor treatment^7,8^. Thus, it is possible that drug resistant cancer cells evade RasG12C inhibitors by rewiring their growth signaling programs to rely on WT H/NRas. Notably, one of the major differences between KRas and H/NRas is their subcellular localization. While all Ras isoforms are localized to the PM, KRas4A can localize to the mitochondria and HRas and NRas isoforms dynamically shuttle between the PM, endoplasmic reticulum (ER), and golgi^10,11^. Thus, it is possible that RasG12C inhibition leads to spatial re-organization (more to endomembrane regions) of Ras activities to rely more on WT H/NRas signaling to permit drug resistance. If true, increased Ras activities at endomembranes instead of PM may enable new protein-protein and enzyme-substrate interactions that turn on different signaling programs.

To probe this hypothesis, we used our recently developed genetically encoded sensors to measure endogenous Ras activity (Ras-LOCKR-S (LOCKR: Latching Orthogonal Cage-Key pRotein)) and environment (Ras-LOCKR-PL)^12^. To differentiate WT versus mutant Ras activity, Ras-LOCKR tools were localized to particular subcellular regions to measure compartment specific Ras activities. Single-cell analysis of localized Ras activities using our Ras-LOCKR-S demonstrated that a subpopulation of KRasG12C-driven cancer cells treated with RasG12C-GDP inhibitors have increased golgi-localized WT H/NRas activity which is responsible for rebound MAPK signaling and drug resistant cell growth. Application of golgi-localized Ras-LOCKR-PL to understand the components responsible for golgi-localized Ras activity led to the identification of Major Vault Protein (MVP). Pairing structure (MVP interactome) with function (golgi-Ras activity) at a single-cell level demonstrated that the high golgi-Ras activity subpopulation is driven by MVP interacting with several MAPK pathway components (Shp2, Erk, RTKs, Ras). In addition, MVP drives mitochondria-localized Ras activities presumably through KRas4A (the only major Ras isoform there^13^), which interacts with the mitochondrially localized voltage-dependent anion channels (VDAC) and in turn alters metabolite availability. Recently developed RasG12C-GTP inhibitors may provide another avenue to tackle KRasG12C-driven cancer cells^5^, however we saw that RasG12C-GTP inhibition also led to MVP-dependent WT H/NRas activation. Altogether, linking structure to function at a single-cell level by using our Ras-LOCKR tools enabled the discovery of MVP driving a subpopulation of cancer cells to evade RasG12C inhibitors. Thus, this study fills in the gap of our knowledge about the signaling components determining RasG12C inhibitor resistance.

## Results

### AMG-510-promoted Ras activation at the golgi enables rebound MAPK signaling and drug resistant cell growth

To understand mechanisms of resistance against inhibitors targeting RasG12C-GDP (**Figure 1A**), we studied two KRasG12C-expressing cell lines (H358 (heterozygous KRasG12C) and MIA PaCa-2 (homozygous KRasG12C)) that show different MAPK signaling (measured by pErk levels) rebound kinetics following treatment with AMG-510 (**Figure 1B**). As RasG12C-GDP inhibition has been shown to lead to non-uniform changes across cells at the genetic and transcriptomic level^4,6^, we probed whether rebound MAPK signaling downstream of RasG12C is also heterogeneous by measuring the single-cell distribution of phospho-Erk (pErk) levels via immunostaining. In H358 cells, we observed that pErk levels were relatively uniform (coefficient of variation (CV)=8.5%) before AMG-510 treatment but highly heterogenous (CV= 35% (24 hr), 42% (48 hr), and 30% (72 hr)) at timepoints where rebound signaling was observed (**Figure 1C**). MIA PaCa-2s showed similarly heterogenous pERK levels in response to AMG-510 treatment (**Figure S1A**). Thus, RasG12C-GDP inhibition leads to a non-uniform adaptive signaling response, with a subpopulation of cells demonstrating high pErk levels, which we termed “AMG-510-promoted signaling adaptive cell subpopulation”.

**Figure 1:**
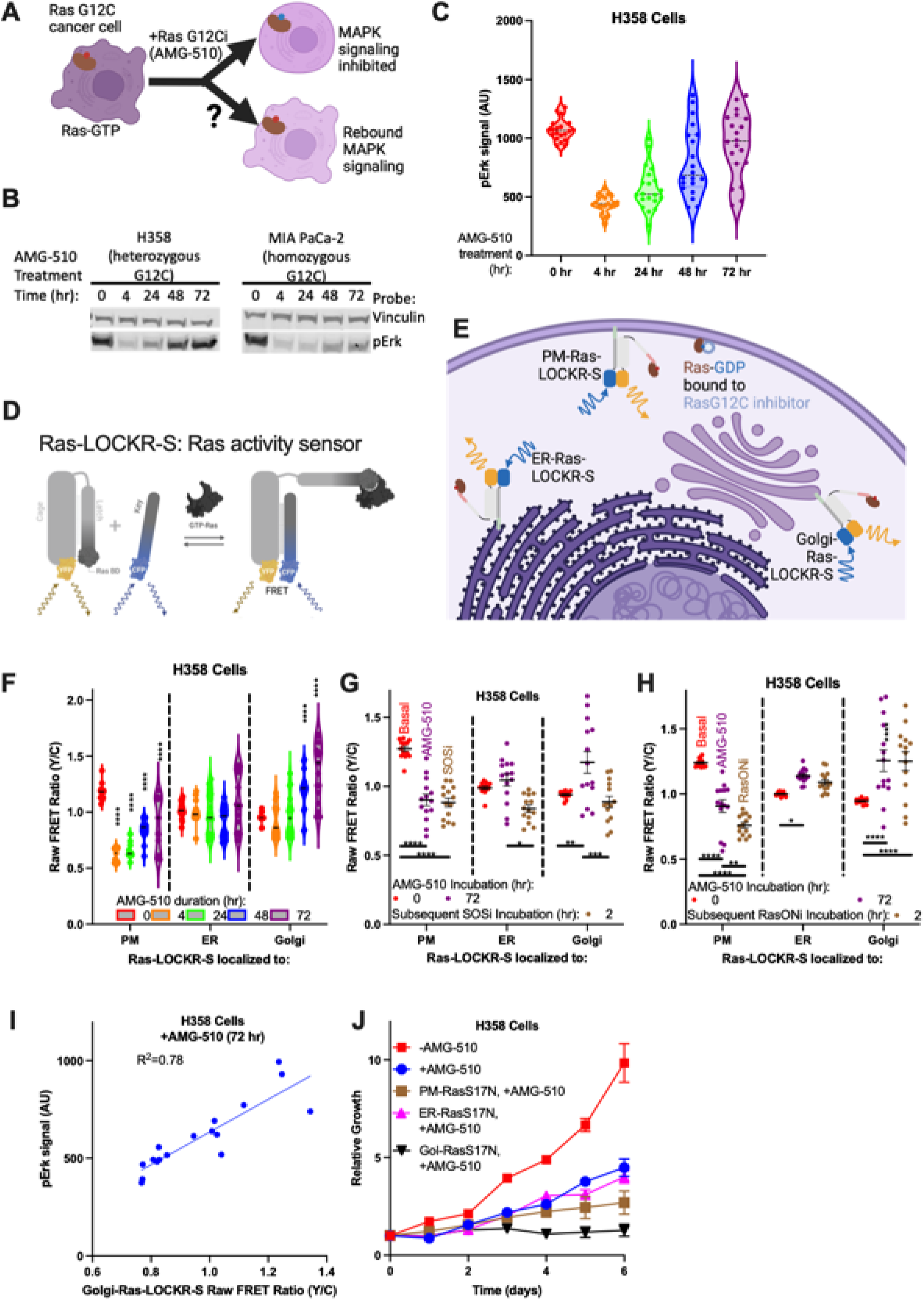
RasG12C-GDP inhibitor (AMG-510) treatment leads to golgi-localized Ras activity that drives oncogenic MAPK signaling. (A) Schematic showing the question being studies: how does RasG12C inhibition lead to adaptive signaling and drug resistance? (B) Representative immunoblot of H358 (lung KRasG12C heterozygous) and MIA PaCa-2 (pancreas KRasG12C homozygous) cells treated for the times indicated with 100nM AMG-510. (C) Violin plots of phospho-Erk (pErk) levels in H358 cells treated with 100nM AMG-510 for the times indicated. Each dot represents the average fluorescence intensity (from pErk immunostaining) of a single cell (n=20 cells per condition). Coefficient of variation (CV) at 0 hour=8.5%, 4 hour=17%, 24 hour=35%, 48 hour=42%, 72 hour=30%. (D) Schematic of Ras-LOCKR-S; a genetically encoded sensor for measuring endogenous Ras activity. (E) Schematic of localized Ras-LOCKR-S at three subcellular regions. (F) Raw FRET ratios (Y/C) of localized Ras-LOCKR-S in H358 cells treated for the times indicated with 100nM AMG-510 (n=17 cells per condition). Statistics: Ordinary two-way ANOVA. PM: CV at 0 hour=5.7%, 4 hour=10%, 24 hour=11%, 48 hour=11%, 72 hour=26%. ER: CV at 0 hour=7%, 4 hour=9.4%, 24 hour=19%, 48 hour=15%, 72 hour=23%. Golgi: CV at 0 hour=4.6%, 4 hour=16%, 24 hour=19%, 48 hour=15%, 72 hour=19% (G-H) Raw FRET ratios (Y/C) of localized Ras-LOCKR-S in H358 cells treated 72 hours with 100nM AMG-510 (n=17 cells per condition). After 72 hour AMG-510 treatment, either 1μM Sos inhibitor BI-3406 (G) or 100nM RasG12C-GTP (RasON) inhibitor (RasONi) RMC-6291 (H) were added for 2 hours. Statistics: Ordinary two-way ANOVA. (I) Scatterplot comparing golgi-Ras-LOCKR-S raw FRET ratio to pErk immunostaining fluorescence in H358 cells treated with 100nM AMG-510 for 72 hours. Each dot represents a single cell where both measurements were done (n=17 cells per condition). (J) Cell growth curves of H358 cells expressing localized RasS17N and treated with DMSO or 100nM AMG-510 (n=3 experiments per condition). Each data point represents the average of 3 experiments.

As pErk levels non-uniformly rebound following AMG-510 treatment (**Figure 1C and S1A**), we next determined if these differences in downstream signaling correlate to heterogeneity in Ras activation state (Ras-GTP). To do this, we used a genetically encoded Ras sensor (Ras-LOCKR-S) that we recently developed^12^, which allows the measurement of endogenous Ras activity at a single-cell level with subcellular resolution (**Figure 1D**). Ras-LOCKR-S uses two *de novo* switchable proteins (“Cage” and “Key”)^14–16^ fused to fluorescent proteins (FPs), which allow Ras-GTP levels to be measured as increases in Förster Resonance Energy Transfer (FRET, measured by FRET ratio=FRET channel/donor FP channel) (**Figure 1D**).

Instead of determining total levels of Ras activation in each cell, we measured subcellular Ras activities because this would allow us to differentiate between reactivation of mutant KRasG12C, which is mainly localized to the plasma membrane, from activation of wildtype (WT) HRas and NRas (H/NRas), which are enriched at endomembranes such as the ER and golgi^17,18^. To measure subcellular Ras activities, the Key of Ras-LOCKR-S was localized to either the PM, ER, or golgi by fusing N-terminal localization sequences (**Figure 1E and S1B**). In KRasG12C cells, AMG-510 addition led to rapid (4 hr) decreases followed by a gradual, partial rebound in PM-Ras-LOCKR-S FRET ratios (**Figure 1F and S1C**). In contrast, AMG-510 treatment led to increased levels of Ras activation at endomembranes. H358 cells displayed no significant changes in ER Ras activity but robust increases in golgi-Ras-LOCKR-S FRET ratios (**Figure 1F**) and MIA PaCa-2s showed FRET ratio increases with both ER-Ras-LOCKR-S and golgi-Ras-LOCKR-S (**Figure 1F and S1C**). Similar to the non-uniform pErk levels we observed (**Figure 1C**), AMG-510 treatment resulted in heterogenous (CV at 72 hour=19%) increases in Ras-GTP levels at endomembranes in both KRasG12C cells, especially in rebound-prone H358 cells (**Figure 1F and S1C**). Altogether, these results demonstrate that AMG-510 treatment causes Ras activation at endomembranes in a subpopulation of cells possibly to compensate for inhibition of KRasG12C-GDP.

We reasoned that AMG-510-promoted Ras activation at endomembranes is a result of WT H/NRas activation at the ER and golgi due to the subcellular distributions of these Ras isoforms. To test whether WT H/NRas plays a role in redistributed Ras signaling in response to Ras G12C inhibition, we measured how subcellular Ras activation levels respond to inhibition of Sos1 GEF because WT H/NRas show a greater dependence on GEF activity than KRasG12C. Rebound-prone H358 cells transiently expressing subcellularly localized Ras-LOCKR-S were treated with AMG-510 for 72 hours followed by treatment with the Sos1 inhibitor BI-3406 (SOSi, BI-3406^19^) for 2 hours (**Figure 1G and S1D**). SOSi treatment significantly decreased AMG-510-promoted golgi-Ras activation but minimally affected Ras-GTP levels at the PM (**Figure 1G and S1D**), suggesting that AMG-510 leads to WT H/NRas activity at the golgi. We also tested the subcellular contribution KRasG12C plays in redistributed Ras signaling in response to AMG-510 treatment with an inhibitor that selectively targets GTP-bound RasG12C (RasONi, RMC-6291^5^). H358 cells transiently expressing PM, ER, or golgi-localized Ras-LOCKR-S were subjected to AMG-510 for 72 hours, and then treated with RasONi for 2 hours (**Figure 1H and S1E**). RasONi treatment significantly decreased AMG-510-promoted PM-Ras but not golgi-Ras activation (**Figure 1H and S1E**), suggesting that AMG-510-insensitive KRasG12C-GTP is present at the PM possibly by its overexpression as a drug adaptation. Altogether, these results indicate that WT H/NRas is a major contributor for redistributed signaling at the golgi while GTP-bound KRasG12C plays a role for redistributed PM-localized signaling.

While each of the three subcellular regions tested (PM, ER, and golgi) showed dynamic changes in Ras activities after AMG-510 treatment (**Figure 1F and S1C**), we wondered which subcellular pool of active Ras is most relevant to the AMG-510-promoted signaling adaptive cell subpopulation (**Figure 1C**). At the single-cell level, only golgi-Ras activity (measured by golgi-Ras-LOCKR-S FRET ratios) correlated (R^2^=0.78) with Erk activation (pErk immunostaining) 72 hours after AMG-510 treatment in of H358 cells (**Figure 1I and S1G**), suggesting that golgi-localized Ras activity contributes to the AMG-510-promoted signaling adaptive cell subpopulation. Consistent with Ras activity at the golgi playing a functional role in AMG-510-resistant cell growth, only a golgi-localized dominant negative Ras mutant (RasS17N)^20^, which prevented Ras activation at the golgi but not the ER or PM (**Figure S1H**), prevented cell proliferation in KRasG12C cells after AMG-510 treatment (**Figure 1J and S1I**). Golgi-localized RasS17N also prevented global Erk reactivation after 72 hours of AMG-510 treatment (**Figure S1J**). Altogether, we demonstrate that golgi-localized WT H/NRas-GTP drives the rebound MAPK signaling and oncogenic cell growth in the AMG-510-promoted signaling adaptive cell subpopulation. Overall, these results indicate that WT H/NRas plays a critical role in evading KRas G12C inhibition, consistent with previous reports^8,21^.

### Major Vault Protein (MVP) drives local and global changes in signaling during AMG-510 treatment

Our results suggest that WT H/NRas activity at the golgi drives rebound MAPK signaling following Ras G12C-GDP inhibition. To investigate the mechanisms for this activation of Ras, we used another sensor we recently developed^12^ to measure the signaling environment of active Ras (**Figure 2A**). Ras-LOCKR-PL possesses a similar architecture as Ras-LOCKR-S but instead facilitates the profiling of the interactome of active Ras (termed “signalosome”) via assembly of a Ras activity-dependent proximity labeler (**Figure 2A)**. To profile the Ras signalosome at the golgi, we localized the Key of Ras-LOCKR-PL to the golgi (golgi-Ras-LOCKR-PL) (**Figure 2A and S2A**). H358 cells transiently expressing golgi-Ras-LOCKR-PL were co-treated with biotin and either AMG-510 or DMSO for either 4 or 24 hours. Cells were then lysed and biotinylated proteins were enriched with streptavidin beads, followed by tryptic digestion and label-free quantification with mass spectrometry (MS). AMG-510 treatment for 24 hours led to greater protein enrichment compared to DMSO treatment in H358 and MIA PaCa-2 cells (**Figure S2B-C**), consistent with golgi-Ras-LOCKR-S data demonstrating that Ras-GTP levels at the golgi increase after AMG-510 treatment (**Figure 1F and S1C**). We compared the enrichment values (AMG-510/DMSO treatment for given incubation time) of labeled proteins between 4 and 24 hour AMG-510 treatment (labeled “Difference” in **Figure 2B, S2B-C, and Table S1**) as hits more enriched after 24 hour treatment may contribute to the beginning of rebound MAPK signaling (**Figure 2B**). The top enriched proteins (PAPOLG, MVP, and VDAC1) in this “Difference” metric (**Figure 2B**) were verified for their in cell biotinylation with a proximity ligation assay (PLA). In this experiment, we used PLA to probe the co-localization between biotinylated proteins and a candidate protein of interest (POI) (i.e. biotinylated POI), which are detected as fluorescent puncta within cells. Our PLA data confirmed that AMG-510 treatment increased biotinylation of PAPOLG, VDAC1, and MVP in H358 cells (**Figure 2C and S2D**), corroborating the data from our MS profiling experiments (**Figure 2B**). Of these three golgi-Ras signalosome components, only MVP knockdown abrogated AMG-510- promoted golgi-Ras activity in H358 cells (**Figure 2D and S2E-F**), suggesting that MVP plays a role in enabling AMG-510-promoted golgi-Ras activity.

**Figure 2:**
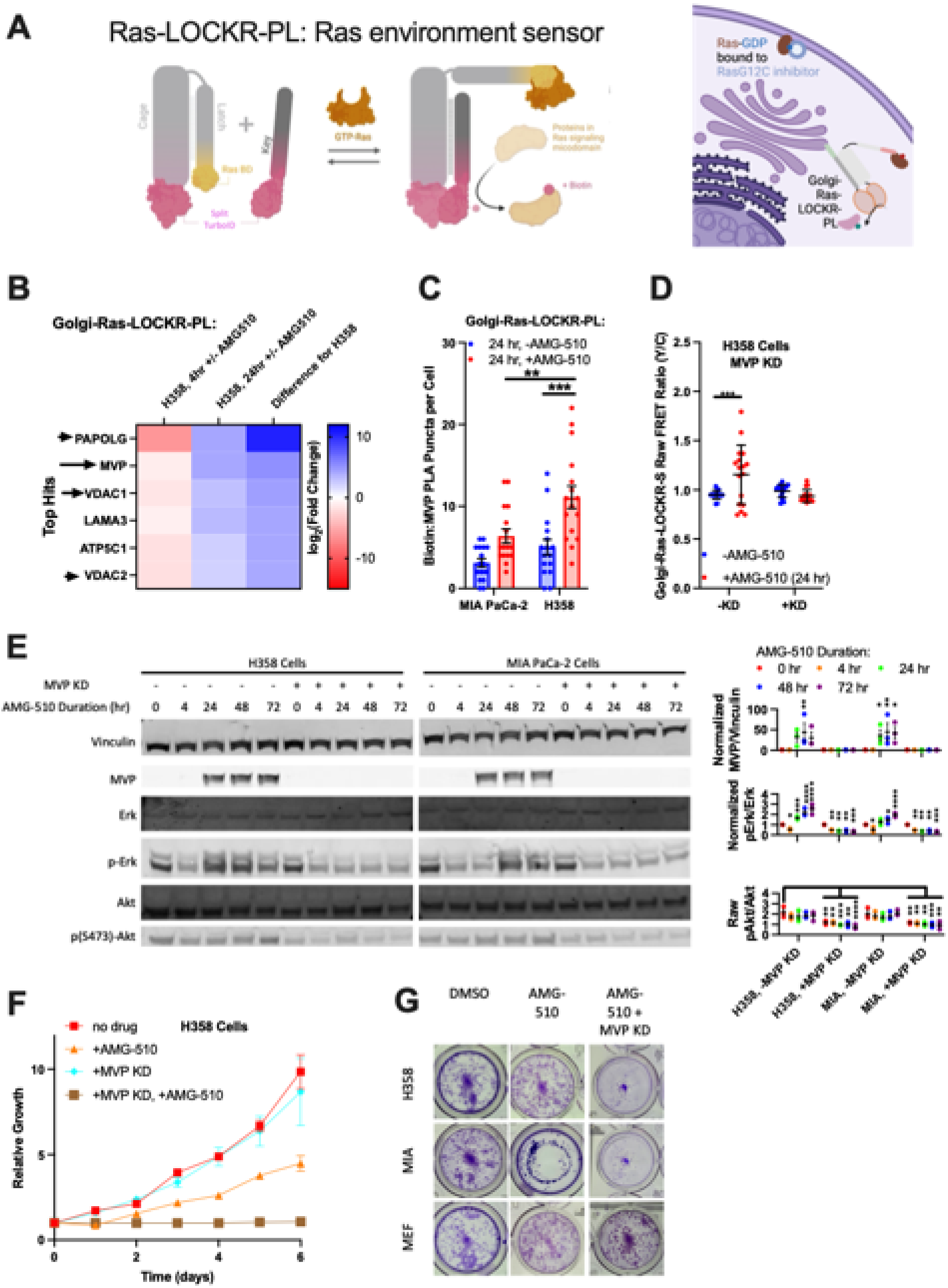
MVP contributes to AMG-510-promoted Ras signaling at the golgi. (A) Left: Schematic of Ras-LOCKR-PL; a Ras activity-dependent proximity labeler that allows the profiling of the environment of active Ras. Right: Schematic of golgi-localized Ras-LOCKR-PL in cells treated with inhibitor for GDP-loaded Ras^G12C^. (B) Heat map of proteins that showed the largest differences in quantification (labeled “Difference”) and -log_2_(fold change) or the difference between 24 and 4 hr AMG-510 treatment. H358 cells expressing golgi-localized Ras-LOCKR-PL were co-treated with 500µM biotin and either 100nM AMG-510 or DMSO followed by lysis, and enrichment with streptavidin beads. Cells were then lysed, trypsinized, and ran through MS (n=4 experimental repeats). With experimental comparisons listed on top, -log_2_(fold change) indicates propensity of protein to be more enriched in the golgi-Ras signaling microdomain when AMG-510 is added. Difference between 24 and 4 hour represents enrichment of proteins in the golgi-Ras signaling compartment throughout time during AMG-510 treatment. Black arrows are MS hits that were tested further throughout this paper. (C) Number of PLA puncta (anti-biotin, anti-MVP) (represents MVP biotinylated by golgi-Ras-LOCKR-PL) per cell. MIA PaCa-2 or H358 cells were transfected with golgi-localized Ras-LOCKR-PL, incubated with 500µM biotin and either with DMSO or 100nM AMG-510, followed by PLA immunostaining (n=16 cells). Each dot represents a single cell and bars represent averages. Statistics: Ordinary two-way ANOVA. (D) Comparison of raw FRET ratios of golgi-Ras-LOCKR-S between MVP KD versus scrambled siRNA in H358 cells treated with 100nM AMG-510 over indicated times (n=17 cells per condition). Statistics: Student’s t-test. (E) Left: Representative immunoblot of H358 and MIA PaCa-2 with or without MVP siRNA transfection and treated over time with 100nM AMG-510. Right: Densitometry quantification of immunoblots either not normalized (pAkt/Akt) or normalized to 0 hr (the rest). Statistics: Ordinary one-way ANOVA, comparison is either indicated or with 0 hour AMG-510 time point. (F) Cell growth curves of H358 cells with MVP KD and/or treatment of 100nM AMG-510 (n=3 experiments per condition). Line represents average from all 3 experiments. (G) Representative images of the colony formation assay of either KRasG12C cells (H358 and MIA PaCa-2) or WT cells (MEF). These cells were treated either with DMSO, 100nM AMG-510 alone, or AMG-510 + MVP KD. Cells were then grown for 2 weeks and then stained by crystal violet.

MVP is a ubiquitous protein that forms a ∼100-mer homo-oligomer structure and its expression has been shown to increase during chemotherapy and radiotherapy^22^. We observed that enrichment of MVP in the golgi-Ras signalosome may possibly be due to MVP shifting its localization to the golgi (**Figure S2G**). Immunoblotting revealed that MVP levels also rapidly (within 24 hours) and significantly increased in KRasG12C cells subjected to AMG-510 (**Figure 2E**), which is consistent with a previous report that also saw statistically significant increases in MVP levels after 24 hour of AMG-510 treatment in MIA PaCa-2 cells^7^. Functionally, MVP KD completely abrogated AMG-510-resistant cell growth and colony formation in KRasG12C cells (**Figure 2F-G and S2H**). MVP has been demonstrated to regulate several signaling branches downstream of Ras by interacting with Shp2 and Erk, nuclearly sequestering negative regulators of PI3K/Akt signaling and thus activating Akt^23^, and promoting RTK-mediated MAPK signaling^24^. However, the role and mechanisms of MVP in terms of drug resistance has not been studied. While our data suggest that MVP plays a local role in golgi-Ras activity, we explored MVP’s role in global signaling. MVP KD eliminated rebound pErk in response to AMG-510 treatment and led to overall decreased pAkt levels (**Figure 2E**). Altogether, these data showcase the role MVP plays in terms of local (golgi-Ras) and global (rebound MAPK, Akt) adaptive signaling in response to RasG12C-GDP inhibiton.

### MVP fuels the AMG-510-promoted signaling adaptive cell subpopulation

Given the effect MVP has on golgi-Ras activity, we wondered if MVP’s presence in the golgi-Ras signalosome can explain the AMG-510-promoted signaling adaptive cell subpopulation. Thus, we used golgi-localized Ras-LOCKR-PL and Ras-LOCKR-S to link structure (Ras signalosome) to function (Ras activity), respectively, at the single-cell level. Both MVP’s proximity to the golgi-Ras signalosome and golgi-Ras activity were measured in KRasG12C cells transiently expressing golgi-Ras-LOCKR-S and golgi-Ras-LOCKR-PL and subjected to DMSO or AMG-510 for 24 hours (**Figure 3A-B**). Similar to pErk levels (**Figure 1C**), both golgi-Ras-LOCKR-PL-mediated biotinylation of MVP (measured by PLA) and golgi-Ras-LOCKR-S FRET ratios were heterogeneous in response to AMG-510 treatment (**Figure 3B**). This labeling heterogeneity reflected the relative strength of golgi-Ras activity as evidenced by the positive correlation (R^2^=0.66) between golgi-Ras activity and golgi-Ras-LOCKR-PL-mediated MVP biotinylation in KRasG12C cells (**Figure 3C**). Satisfyingly, this single-cell analysis made possible by Ras-LOCKR tools demonstrated that the subpopulation of cells with high golgi-Ras activity (AMG-510-promoted signaling adaptive cell subpopulation) are typified by the presence of MVP in the golgi-Ras signalosome but not by MVP expression levels (**Figure S3A**). Further confirming MVP’s functional role in AMG-510-mediated Ras activation at the golgi, MVP KD suppressed golgi-Ras activity 72 hours after AMG-510 treatment, suggesting that MVP plays a strong role in WT H/NRas activation at the golgi in response to RasG12C-GDP inhibition (**Figure 3D and S3B-D**). Thus, these results reveal MVP as an upstream driver of golgi-localized WT H/NRas activity that fuels the AMG-510-promoted signaling adaptive cell subpopulation.

**Figure 3:**
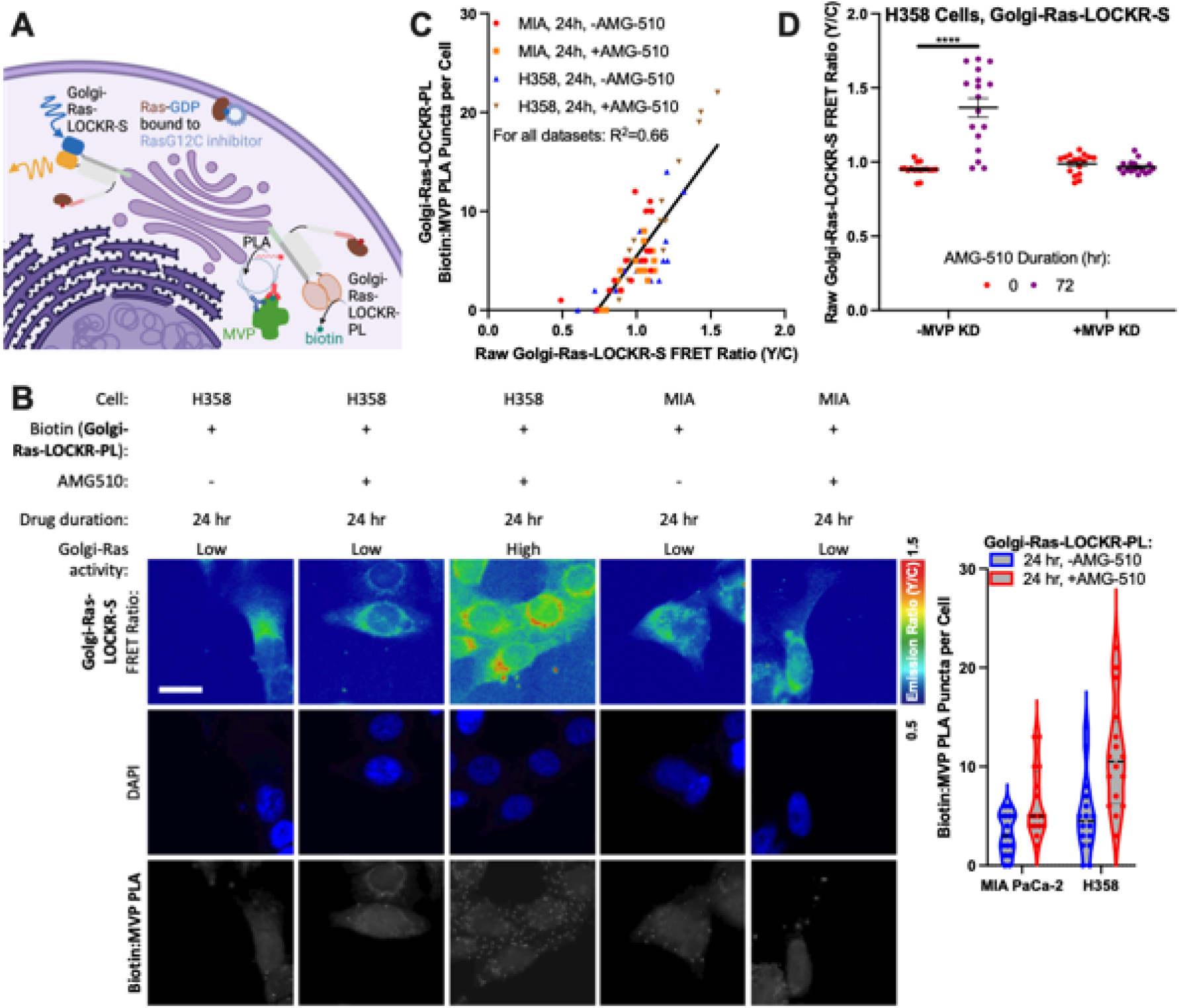
MVP at the golgi fuels the Amg-510-promoted signaling adaptive cell subpopulation. (A) Schematic of the single-cell structure-function relationship mapping experiment. golgi-Ras-LOCKR-PL with PLA and golgi-Ras-LOCKR-S were used to relate MVP’s presence in the golgi-Ras signalosome (structure) to golgi-Ras activity (function) in response to AMG-510 treatment. (B) Right: Representative epifluorescence images of H358 or MIA PaCa-2 cells expressing golgi-Ras-LOCKR-PL and golgi-Ras-LOCKR-S. Cells were co-treated with 500µM biotin and either DMSO or 100nM AMG-510 for 24 hours. Cells were then imaged and subjected to PLA using anti-biotin and anti-MVP antibodies. Scale bar = 10µm. Left: Violin plot comparing PLA puncta (anti-biotin, anti-MVP) per cell. MIA PaCa-2 or H358 cells were transfected with golgi-localized Ras-LOCKR-PL, incubated with 500µM biotin and DMSO or 100nM AMG-510 100nM AMG-510, and underwent PLA immunostaining (n=16 cells). Each dot represents a single cell. Same dataset is used in Fig. 2C. Statistics: CV for MIA PaCa-2 -AMG-510=66%, CV for MIA PaCa-2 +AMG-510=77%, CV for H358 -AMG-510=54%, CV for H358 +AMG-510=50%. (C) Scatterplot comparing golgi-Ras-LOCKR-PL-mediated labeling of MVP (y-axis) to golgi-Ras-LOCKR-S raw FRET ratios (x-axis). Each dot represents a single cell where both measurements were performed (n=16 cells per condition). (D) Raw FRET ratios of golgi-Ras-LOCKR-S in H358 cells with MVP KD or scrambled siRNA and then treated for 72 hours with 100nM AMG-510 (n=17 cells per condition). Statistics: Student’s t-test.

### MVP scaffolds MAPK pathway components to enable oncogenic signaling

We next investigated the molecular details in how MVP can activate Ras and its downstream MAPK pathway during AMG-510 treatment. It has been shown that MVP becomes tyrosine phosphorylated and binds to Shp2 and Erk upon EGFR stimulation^24^. Given the link between MVP and MAPK signaling, we investigated MVP’s interactome (structure) with MAPK pathway components (Shp2, Erk, RTKs EGFR and fibroblast growth factor receptor (FGFR)) to golgi-Ras activities (function) at the single-cell level. Using PLA, H358 cells treated with AMG-510 for 24 hours displayed significant increases in MVP’s proximity to Shp2, Erk, EGFR, and FGFR3 (**Figure 4A**). We performed single-cell structure-function analysis and saw MVP’s proximity to Shp2, Erk, EGFR and FGFR3 positively correlated with golgi-Ras-LOCKR-S FRET ratios (R^2^=0.85) (**Figure 4B**), suggesting that increased MVP complex formation with MAPK signaling components enhances golgi-Ras activity. Moreover, MVP KD decreased Shp2:Erk and Shp2:EGFR complexes that were promoted by AMG-510 treatment in KRasG12C cells (**Figure 4C**). At the single-cell level, Shp2’s proximity to Erk and EGFR positively correlated (R^2^=0.83) with golgi-Ras-LOCKR-S FRET ratios (**Figure 4D**), suggesting that MVP acts as an intermediate scaffold for Shp2 to interact with its substrates. Given the connection of MVP to WT H/NRas activity at the golgi (**Figure 3D**), we wondered if MVP interacts with Ras itself. Using the best available antibodies for the different Ras isoforms^25^, we observed that MVP’s interactions with WT HRas and NRas increased upon AMG-510 treatment but not for KRas (**Figure S4A-B**). At the single-cell level, MVP’s complex formation with HRas and NRas, as measured by PLA, correlated well with golgi-Ras-LOCKR-S FRET but mutant KRas’s proximity to MVP negatively correlated with golgi-Ras activity (**Figure S4C**), presumably due to HRas and NRas being endomembrane localized. These results suggest that MVP forms a complex with WT H/NRas to promote adaptive MAPK signaling in response to RasG12C-GDP inhibition.

**Figure 4:**
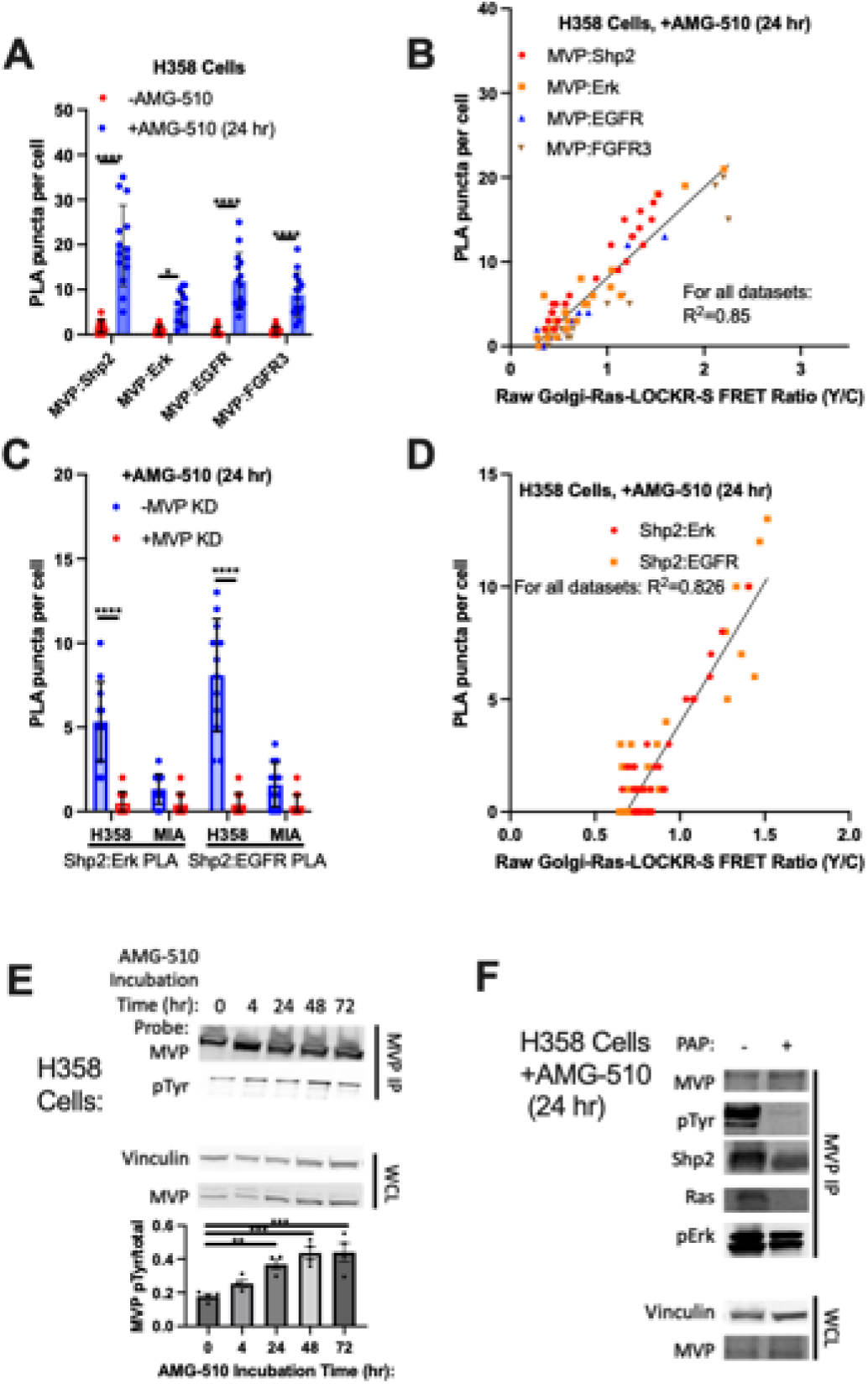
Structure-function mapping shows that MVP interacts with MAPK pathway components to drive the Amg-510-promoted signaling adaptive cell subpopulation. (A-B) H358 or MIA PaCa-2 cells transfected with golgi-Ras-LOCKR-S and treated with DMSO or 100nM AMG-510 for 24 hours, followed by probing for MVP/ MAPK pathway component proximity with PLA. Displayed is (A) quantification across conditions and (B) correlation between golgi-Ras-LOCKR-S raw FRET ratios and PLA puncta (anti-MVP, anti-MAPK pathway component) per cell (n=14 cells per condition). Same dataset is used in Fig. 4A. Statistics: Student’s t-test. (C-D) H358 or MIA PaCa-2 cells transfected with golgi-Ras-LOCKR-S, transfected with MVP or scrambled siRNA for 2 days, incubated with 100nM AMG-510 for 24 hours, and immunostained and probed for Shp2:Erk and SHP2:EGFR proximity via PLA. Displayed is (C) quantification across conditions and (D) correlation between golgi-Ras-LOCKR-S FRET ratios and PLA puncta (anti-Shp2, anti-Erk, anti-EGFR) per cell (n=12 cells per condition). Same dataset is used in Fig. 4C. Statistics: Student’s t-test. (E) H358 cells were treated with 100nM AMG-510 for 72 hours. Samples were then lysed, MVP was immunoprecipitated (IPed), and total MVP and pTyr levels were determined via immunoblotting. Top: Representative immunoblot with IP samples on top and whole cell lysate (WCL) samples on bottom. Bottom: Densitometry quantification of immunoblots where the ratio between MVP with phospho-tyrosine modification (pTyr in the MVP IP) and MVP pulled down (n=4 biological replicates). Statistics: Student’s t-test. (F) H358 cells were treated with 100nM AMG-510 for 24 hours. Samples were then lysed, treated with or without potato acid phosphatase (PAP) for 45 minutes, MVP was IPed, and levels of Shp2, Ras, and pERK that were co-IPed were measured using immunoblotting. Representative immunoblot with IP samples on top and WCL samples on bottom (n=3 biological replicates)

We next investigated how MVP interacts with MAPK signaling components. A previous study showed that EGFR stimulation leads to phosphorylation of MVP tyrosines, which leads to Shp2 recruitment through its SH2 domain.^24^ Thus, we probed whether MVP becomes tyrosine phosphorylated in response to AMG-510 treatment^26^. Indeed, we found that AMG-510 treatment led to a steady increase in MVP tyrosine phosphorylation (**Figure 4E**). To test whether tyrosine phosphorylation is necessary for MVP’s interaction with MAPK pathway components, H358 cells were treated with AMG-510 for 24 hours and lysate was then subjected to potato acid tyrosine phosphatase (PAP) treatment^27^, followed by MVP immunoprecipitation, and immunobloting. PAP treatment led to decreased Shp2, Ras (pan-Ras), and pErk co-immunoprecipitation with MVP (**Figure 4F**), demonstrating that MVP’s interaction with MAPK components is mediated through tyrosine phosphorylation.

Altogether, by pairing localized Ras signalosome profiling (Ras-LOCKR-PL) with subcellular Ras activities (Ras-LOCKR-S), we were able to link structure to function to understand how the AMG-510-promoted signaling adaptive cell subpopulation can arise. Our data suggests that by bringing substrates and clients of the same pathway together, MVP scaffolding of MAPK components (Shp2, Erk, RTKs, WT H/NRas) enhances golgi-Ras activity, which promotes adapative oncogenic signaling.

### AMG-510 treatment alters cytosolic ATP/ADP ratios and mitochondrial Ras activity

Interestingly, mitochondrial VDAC1 and VDAC2 are two of the most prominently labeled proteins by golgi-Ras-LOCKR-PL in response to AMG-510 treatment (**Figure 2B**), possibly due to golgi-mitochondria contacts^28,29^ (**Figure S5A**). VDACs are outer mitochondrial membrane channel proteins that regulate cytosolic and mitochondrial levels of ATP, which affects glycolytic rates based on cytosolic ATP’s ability to inhibit several glycolytic enzymes^30^. KRas4A directly enhances glycolytic flux by blocking allosteric inhibition of hexokinase^31^. Recent reports indicate that RasG12C-GDP inhibition leads to increased mRNA and protein expression of glycolytic enzymes with a concomitant increase in the levels of several intermediate metabolites of glycolysis^7,9^. Consistent with these findings, we observed that treating RasG12C-GDP-expressing cells with AMG-510 for 72 hour led to increased glucose consumption and lactate release, signatures of enhanced glycolytic flux (**Figure 5A**). To determine whether VDAC function is altered in response to AMG-510 treatment as a possible mechanistic link to the observed glycolytic changes (**Figure 5A**), we next used the fluorescence-based biosensor Cyto-PercevalHR^32,33^, which allows the measurement of cytosolic ATP/ADP ratios (500nm/450nm fluorescence emission ratios) at the single-cell level. We observed a heterogenous (CV at 72 hour=28%) reduction in cytosolic ATP/ADP ratios (**Figure 5B and S5B-C**) in response to AMG-510 treatment, suggesting that RasG12C-GDP inhibition alters VDAC’s function. Thus, RasG12C-GDP inhibition leads to a “metabolically adaptive subpopulation” akin to the signaling adaptive cell subpopulation (**Figure 1**).

**Figure 5:**
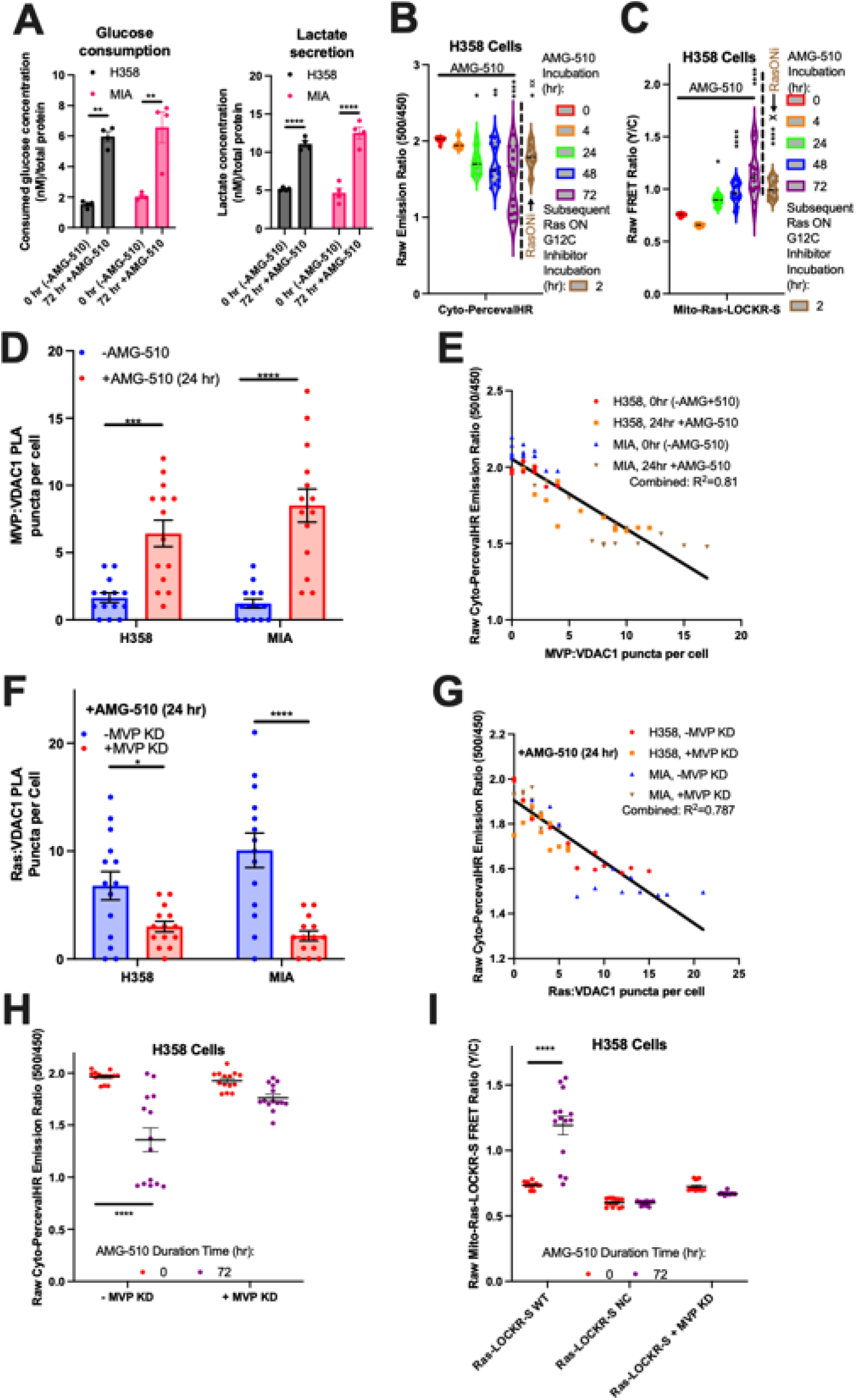
MVP is required for the metabolically adapting cell subpopulation promoted by AMG-510. (A) H358 and MIA PaCa-2 cells were measured for glucose consumption and lactate release before or 72 hours after 100nM AMG-510 addition (n=4 experiments per condition) (B-C) H358 cells transfected with either cyto-PercevalHR (B) or mito-Ras-LOCKR-S (C) were first treated with 100nM AMG-510 for the times indicated followed by incubation with 100nM RasONi for 2 hours (n=14 cells per condition). Shown are either the raw emission ratios (500/450) from cyto-PercevalHR (B) or raw FRET ratios from mito-Ras-LOCKR-S (C). Statistics: Ordinary two-way ANOVA. * is comparison to 0 hour AMG-510 time point and x is comparison to 72 hour AMG-510 time point. CytoPercevalHR: CV at 0 hour=1.5%, CV at 4 hour=3.4%, CV at 24 hour=9.6%, CV at 48 hour=15%, CV at 72 hour=28%, CV after RasONi=9.4%. mito-Ras-LOCKR-S: CV at 0 hour=1.6%, CV at 4 hour=2%, CV at 24 hour=5.2%, CV at 48 hour=9.8%, CV at 72 hour=20%, CV after RasONi=8.8%. (D-E) H358 or MIA PaCa-2 cells transfected with cyto-PercevalHR, incubated with DMSO or 100nM AMG-510 for 24 hours, imaged, and then underwent PLA (anti-MVP, anti-VDAC1). Displayed is (D) quantification across conditions and (E) correlation between cyto-PercevalHR emission ratios and PLA puncta (anti-MVP, anti-VDAC1) per cell (n=14 cells per condition). Statistics: Student’s t-test. (F-G) H358 or MIA PaCa-2 cells transfected with Cyto-PercevalHR, transfected with scrambled siRNA or MVP siRNA, treated with 100nM AMG-510 for 24 hours, imaged, and then underwent PLA (anti-pan Ras, anti-VDAC1). Displayed is (F) quantification across conditions and (G) correlation between cyto-PercevalHR emission ratios and PLA puncta (anti-pan Ras, anti-VDAC1) per cell (n=14 cells per condition). Statistics: Student’s t-test. (H) Raw emission ratios of H358 cells transfected with cyto-PercevalHR and either with scrambled siRNA or MVP siRNA and treated for 72 hours with 100nM AMG-510 (n=14 cells per condition). Statistics: Student’s t-test. (I) Raw FRET ratios of H358 cells transfected with either mito-Ras-LOCKR-S WT, mito-Ras-LOCKR-S NC sensor, or mito-Ras-LOCKR-S WT and MVP siRNA, and treated for 72 hours with 100nM AMG-510 (n=14 cells per condition). Statistics: Student’s t-test.

We suspected that the AMG-510-promoted upregulation of glycolysis (**Figure 5A**) is due to AMG-510 modulating the KRas4A spliceform. KRas4A is the only major Ras isoform localized at the outer mitochondrial membrane^13^, thus as a proxy for mitochondrial Ras (mito-Ras) activities we measured mitochondrial Ras-GTP levels by localizing the Key of Ras-LOCKR-S to the outer mitochondrial membrane (**Figure S5D**). AMG-510 treatment led to an initial decrease (4 hours) followed by a gradual heterogenous (CV at 72 hour=20%) increase in mito-Ras-LOCKR-S FRET ratios that exceeds basal levels (**Figure 5C and S5E-F**), which we interpret as an initial block of basal KRas4A activity but then a significant increase in KRas4A-GTP levels, possibly due to adaptive overexpression of mutant KRas^6^. As a control, no FRET ratio increases were seen in a negative control (NC) mito-Ras-LOCKR-S sensor (**Figure S5E, F**) that possesses a mutant RasBD incapable of binding Ras-GTP^12^. Unlike golgi-Ras activity, mito-Ras activity does not correlate with global pErk levels (**Figure S5G**), suggesting that mito-Ras activity is not responsible for AMG-510-promoted signaling adaptive cell subpopulation. To test whether AMG-510-promoted increases in cytosolic ATP/ADP ratios and mitochondrial Ras-GTP are the result of activating WT or mutant Ras, RasONi was added 72 hour after AMG-510 treatment. Subsequent RasONi treatment for 2 hours partially (∼50%) reversed AMG-510-promoted increases in mito-Ras activities and decreases in cytosolic ATP/ADP ratios (x in **Figure 5B-C** compares 72 hour AMG-510 alone versus 72 hour AMG-510 followed by 2 hour RasONi), suggesting that mutant KRas4A is likely responsible for the observed mitochondrial Ras activity and metabolic alterations that result from RasG12C-GDP inhibition. Together, AMG-510 leads to changes in mitochondrial function which is reflected in altered cytosolic ATP/ADP ratios and mitochondrial Ras activities, and these correspond to enhanced glycolysis (see Discussion).

### MVP mediates AMG-510-promoted changes in cytosolic ATP/ADP ratios and mito-Ras activities

After establishing an AMG-510-promoted metabolically adaptive cell subpopulation (**Figure 5B**), we were interested in determining what drives this subpopulation. As we previously saw that MVP affects golgi-Ras activity during AMG-510 treatment by interacting with MAPK pathway components (**Figure 4**), we wondered if MVP also plays a role in altering mito-Ras activities and cytosolic ATP/ADP ratios by locally scaffolding signaling effectors and VDAC1, which we observed is within the Ras signalosome in response to AMG-510 treatment (**Figures 2B and S2D**). Hence, we performed single-cell structure-function analysis to relate MVP interactome (structure) to function (changes in cytosolic ATP/ADP ratios). Using PLA to measure protein co-localizations at the single-cell level, we observed that AMG-510 treatment resulted in increased MVP:VDAC1 co-localization after AMG-510 addition (**Figure 5D**) and that these proximities negatively correlated with cytosolic ATP/ADP ratios (R^2^=0.81) (**Figure 5E**). MVP also appears to mediate AMG-510-promoted Ras:VDAC1 complex formation as KD of MVP led to decreased Ras:VDAC1 co-localization as measured by PLA (**Figure 5F**). The level of VDAC1 in proximity to Ras negatively correlated (R^2^=0.79) with cytosolic ATP/ADP ratios (**Figure 5G**), suggesting that MVP-mediated Ras:VDAC1 co-localization fuels the AMG-510-promoted metabolically adaptive subpopulation. We observed that MVP is also necessary for AMG-510-promoted changes in metabolism, as no decrease in cytosolic ATP/ADP ratios were observed upon AMG-510 treatment of MVP KD cells (**Figure 5H and S5B-C**). Consistent with MVP also being required for mitochondrial Ras signaling, MVP KD blunted AMG-510-promoted increases in mito-Ras activities (**Figure 5I and S5E-F**). Overall, our single-cell analyses capture the AMG-510-promoted metabolically adapting subpopulation and reveals that MVP drives this subpopulation by recruiting factors to the mitochondria to regulate KRas4A and VDAC (see Discussion for a potential mechanism), thus altering metabolism.

### Inhibition of RasG12C-GTP (RasONi) also leads to MVP-dependent, golgi-localized Ras activity

The development of new inhibitors to GTP-loaded RasG12C (distinct from AMG-510 which targets RasG12C-GDP) such as RasON inhibitor (RasONi, RMC-6291^5^) lead to more potent inhibition of mutant RasG12C as RasG12C is primarily GTP-loaded and this is the signaling competent form. Inhibitors for RasG12C-GTP also more quickly inhibit RasG12C as RasG12C hydrolysis of GTP is slow, thus in theory should limit the time adaptive signaling can emerge. Moreover, resistance to RasG12C-GDP inhibitors can be mediated by cells overexpressing RasG12C, which preferentially are bound to GTP^6^. For these reasons, RasG12C-GTP inhibitors may be more clinically promising drugs that may be harder for cancer cells to develop resistance against. Unfortunately, when RasONi was applied to KRasG12C cells for over 72 hours, pErk levels initially decreased but later rebounded after 48- and 72-hours (**Figure 6A**) similar to the pErk dynamics after AMG-510 incubation (**Figure 1B**). Similar to AMG-510, RasONi treatment led to a significant reduction in Ras-GTP at the PM up to 72 hours but a substantial and heterogenous (CV at 72 hour=29%) increases in golgi-Ras activity, which is not altered when RasONi was refreshed (**Figure 6B and S6A**). Thus, RasONi appears to promote a signaling adaptive cell subpopulation through WT H/NRas activation similar to AMG-510. We previously saw that AMG-510 treatment led to upregulated MVP expression (**Figure 2E**) and MVP drives WT H/NRas activation at the golgi (**Figure 3D**), thus we wondered whether RasONi leads to a similar effect. In KRasG12C cells, MVP expression increased ∼3.5 fold 72 hours after RasONi treatment (**Figure 6C**). MVP KD abrogated golgi-Ras activation in response to RasONi treatment (**Figure 6D and S6B-C**), suggesting that MVP plays a similar role in RasONi-promoted signaling adaptations as it does for RasG12C-GDP inhibition (AMG-510).

**Figure 6:**
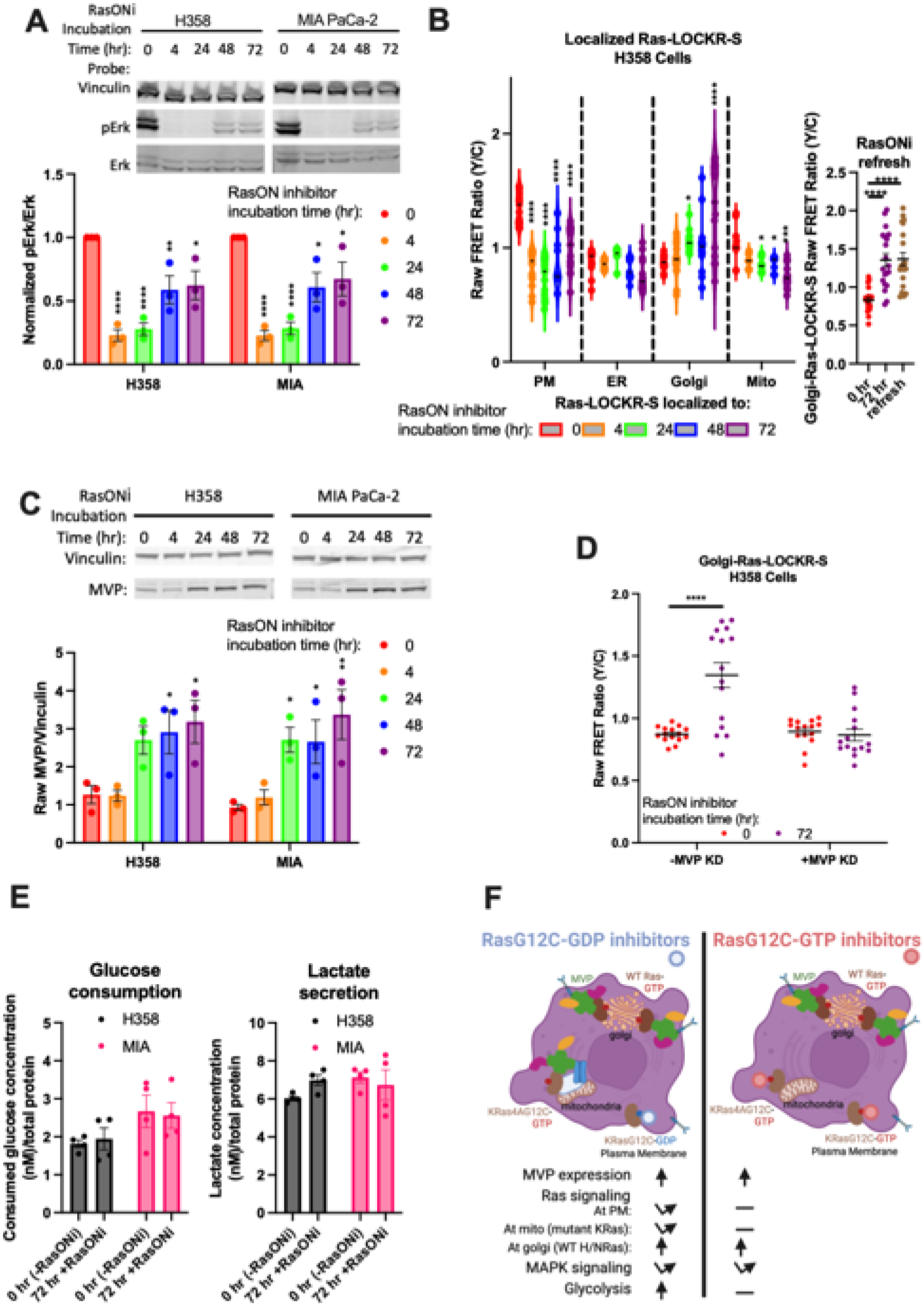
RasG12C-GTP inhibition (RasONi) still leads to rebound oncogenic signaling mediated by MVP. (A) H358 and MIA PaCa-2 cells were treated with 100nM RasONi for the times indicated and then probed for pErk and Erk levels (n=3 experiments). Top: Representative immunoblot. Bottom: Densitometry quantification of immunoblots where pErk over Erk ratios were calculated. Statistics: Ordinary two-way ANOVA, comparison to 0 hour RasONi time point. (B) Left: Raw FRET ratios of H358 cells transfected with subcellularly localized Ras-LOCKR-S and treated for the times indicated with 100nM RasONi (n=15 cells per condition). Statistics: Ordinary two-way ANOVA, comparison to 0 hour RasONi time point. PM: CV at 0 hour=9.9%, 4 hour=19%, 24 hour=24%, 48 hour=27%, 72 hour=19%. ER: CV at 0 hour=11%, 4 hour=4.9%, 24 hour=7.5%, 48 hour=11%, 72 hour=19%. Golgi: CV at 0 hour=7.1%, 4 hour=17%, 24 hour=8.4%, 48 hour=24%, 72 hour=29%. Mito: CV at 0 hour=15%, 4 hour=6.9%, 24 hour=7.3%, 48 hour=8.1%, 72 hour=14%. Right: Raw FRET ratios of H358 cells transfected with golgi-Ras-LOCKR-S and treated over 72 hours with 100nM RasONi and then subsequent 2 hour refresh (labeled “refresh”) of 100nM RasONi. Statistics: Ordinary two-way ANOVA. (C) H358 and MIA PaCa-2 cells treated with 100nM RasONi over 72 hours and probed for MVP and Vinculin (loading control) (n=3 experiments). Top: Representative immunoblot. Bottom: Densitometry quantification of immunoblots where MVP over Vinculin ratios are calculated. Statistics: Ordinary two-way ANOVA, comparison to 0 hour RasONi time point. (D) Raw FRET ratios of H358 cells transfected with either scrambled or MVP siRNA for 2 days, transfected with golgi-Ras-LOCKR-S, and treated for 72 hours with 100nM RasONi (n=15 cells per condition). Statistics: Student’s t-test. (E) H358 and MIA PaCa-2 cells were measured for glucose consumption and lactate release before or 72 hours after 100nM RasONi addition (n=4 experiments per condition). (F) Schematic of MVP scaffolding proteins at the golgi and mitochondria as an adaptive method for resistance to RasG12C inhibitors.

In contrast to AMG-510, RasONi treatment led to prolonged inhibition of PM-Ras-GTP and mito-Ras-GTP (**Figure 6B and S6A**), demonstrating that RasONi can potently inhibit mutant KRas activity. Consistent with RasONi’s inhibition of mito-Ras-GTP, no substantial changes in cytosolic ATP/ADP ratios at the single-cell level (**Figure S6D**) or bulk changes in glycolytic hallmarks (glucose consumption and lactate release, **Figure 6E**) were observed 72 hours after RasONi treatment. Thus, KRasG12C-GTP inhibition results in a clear difference in mitochondrial function relative to KRasG12C-GDP inhibition, most likely do to the ability of RasONi effectively inhibit KRas4A at the mitochondria, thus, limiting metabolic adaptations. Overall, there are similarities and differences between RasG12C-GDP inhibition (AMG-510) versus RasG12C-GTP inhibition (RasONi) (**Figure 6F**). Both inhibitors induced expression of MVP to activate golgi-localized WT H/NRas in order to compensate for mutant KRas inhibition. Distinct between the two inhibitors, mito-Ras activity and glycolysis is promoted only by AMG-510, suggesting that RasONi can more potently inhibit KRas4A and seems to have less effects on metabolism.

In summary, application of our new biosensors and biochemical assays enables us to capture a subpopulation of cells that re-activate oncogenic signaling (MAPK pathway) during both RasG12C-GDP and RasG12C-GTP inhibition. We identified MVP as playing a key role in enabling WT H/NRas activity at the golgi by scaffolding components of MAPK pathway (**Figure 6F**). MVP can also regulate KRas4A activity at the mitochondria to affect metabolic processes (**Figure 6F**). Overall, our work showcases another way cancer cells evade RasG12C inhibition and suggests MVP as a potential important therapeutic target to tackle drug resistance.

## Discussion

Pharmacological targeting of mutant Ras has been a longstanding goal due to its strong oncogenic effect. Only recently have small molecule that are specific and potent enough to provide clinically beneficial inhibition of mutant Ras (RasG12C) been developed. Unfortunately, most patients given RasG12C-GDP inhibitors relapse due to reactivation of the Ras signaling pathway^4^. How cancer cells reactivate the same pathway that was initially inhibited is not completely clear. By analyzing Ras signaling changes at a single-cell level, we aimed to gain mechanistic insight into the molecular signaling components that differentiate drug resistant versus drug responding cell subpopulations.

By using our recently developed activity (Ras-LOCKR-S) and environment (Ras-LOCKR-PL) sensors of endogenous Ras, we detected rebound signaling and rewired metabolism in only a subpopulation of RasG12C cancer cells and identified the mechanisms allowing for this subpopulation to exist. In our study, resistance to RasG12C-GDP inhibitors (AMG-510) is mediated through activation of Ras at endomembranes (golgi and mitochondria) that is dependent on MVP. In addition to Ras signaling, MVP also regulates the activity of Akt during AMG-510 treatment^23^ (**Figure 2E**), which is involved in various cellular processes including metabolism and may provide more insight into RasG12C inhibitor resistance. Recently developed small molecule that specifically inhibit RasG12C-GTP are exciting new modalities for treating RasG12C-driven cancers^5^. However, we observed that RasG12C-GTP inhibition by RasONi also led to rebound MAPK signaling due to MVP-dependent activation of WT H/NRas at the golgi. A previous report^5^ demonstrated that refresh of RasONi in KRasG12C cells led to a decrease in pErk levels, thus the RasONi-promoted WT Ras activation at the golgi may not communicate to productive downstream signaling in this context. Many cases of cancer therapy are associated with increased MVP expression, including previous reports on RasG12C-GDP inhibitors^7^. Our results reveal that Ras-driven cancers exploit MVP to adaptively bypass RasG12C inhibition. More broadly, the MVP-mediated mechanism we observe may be generally involved in resistance to new inhibitors against other Ras mutants^34,35^ and perhaps other cancer drugs.

Mechanistically at the golgi, RasG12C-GDP inhibition promoted WT H/NRas signaling due to MVP-mediated interactions with MAPK pathway proteins (Shp2, Erk, RTKs, Ras). We hypothesize that by recruiting positive regulators to golgi-localized Ras (e.g. Shp2), that either GTP loading is increased or GTP hydrolysis is decreased (or both) to activate WT Ras. However, it is not clear what drives MVP to accumulate at different endomembrane regions such as the golgi (**Figure S2G**). While the structure of oligomeric MVP complexes has been solved^36^, the role of the individual domains has not been carefully investigated. Whether certain domains mediate specific interactions (e.g. with components described here) and/or enable re-localization would be interesting to explore. Another mechanistic question is: how does RasG12C inhibition induce MVP expression? Previous reports show that AMG-510 treatment of *in vitro* RasG12C cell cultures led to increased mRNA^9^ and protein expression^7^ of MVP similar to our results using cultured cell lines. However, whether changes in MVP expression levels occur in cancer patient cells has not been profiled, thus MVP’s clinical relevance is yet still to be explored. MVP expression can be stimulated by cytokines and the immune environment is important for Ras-driven cancers^37^, highlighting the impetus to investigate the role of MVP in more physiologically relevant models. However, the mechanisms underlying MVP expression may not be critical as MVP expression alone does not seem to dictate localized Ras signaling (**Figure S3A**).

At the mitochondria, AMG-510-promoted MVP:VDAC1 co-localization led to decreased cytosolic ATP/ADP ratios and MVP-dependent KRas4A activation, resulting in increased glycolysis. In response to RasG12C-GDP inhibition, we hypothesize that mutant KRas4A at mitochondria is inhibited but then is reactivated with the aid of MVP due to its scaffolding of positive regulators of the MAPK pathway (**Figure 4**). Interestingly, mito-Ras activity does not correlate with pErk levels (**Figure S5G**), which suggests that KRas4A activation at the mitochondria does not productively lead to MAPK signaling which has been shown before^13^. MVP also promotes crosstalk between Ras and VDAC1 (**Figure 5F-G**), thus decreasing cytosolic ATP/ADP ratios (**Figure 5B**). Both KRas4A activation and decreased cytosolic ATP/ADP ratios should enhance glycolytic flux (**Figure 5A**), as KRas4A blocks allosteric inhibition of hexokinase1 and cytosolic ATP inhibits phosphofructokinase and pyruvate kinase^38^. During RasG12C inhibition, MVP regulating metabolism is an intriguing possibility but several mechanistic questions remain. Whether KRas4A activity regulates VDAC1 function or the other way around or are independent events are unclear though. At a molecular level, MVP can associate with KRas (**Figure S4A-C**), mitochondrial KRas4A regulates glycolysis through hexokinase^13^, and VDAC is known to interact with hexokinase^30^. Thus, we hypothesize that a MVP:KRas4A:hexokinase:VDAC multiprotein complex exists. As MVP also recruits other MAPK pathway components such as Erk (**Figure 4A-B and F**) and Erk can phosphorylate and regulate VDAC channel opening^39^, this may be one possible mechanism for MVP-mediated VDAC regulation.

In conclusion, using our new biosensors and biochemical assays, we performed single-cell analysis to capture a subpopulation of RasG12C-addicted cancer cells that evade several RasG12C inhibitors. With these tools, we identified MVP to play a major role in this drug resistance by facilitating protein complexes with MAPK pathway components and metabolite channels. MVP-mediated adaptations drive Ras signaling at endomembranes (golgi and mitochondria), which increases downstream signaling and enables oncogenic metabolism. These results also illustrate how spatiotemporal regulation of central biomolecules such as Ras are necessary for their functional specificity. Overall, this study introduces the technique and application of performing single-cell signaling analyses to understand complex biological phenomena such as drug resistance. By understanding how cancer cells dynamically adapt their signaling, we envision that this analytical methodology will identify additional key factors (e.g. MVP) that enable oncogenesis, targeting of which may have therapeutic benefit.

## STAR Methods

### KEY RESOURCES TABLE

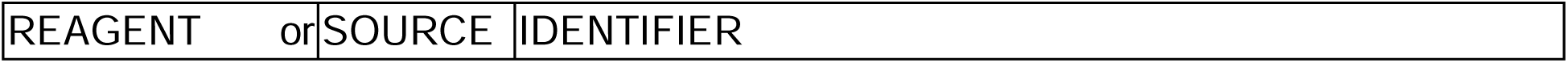
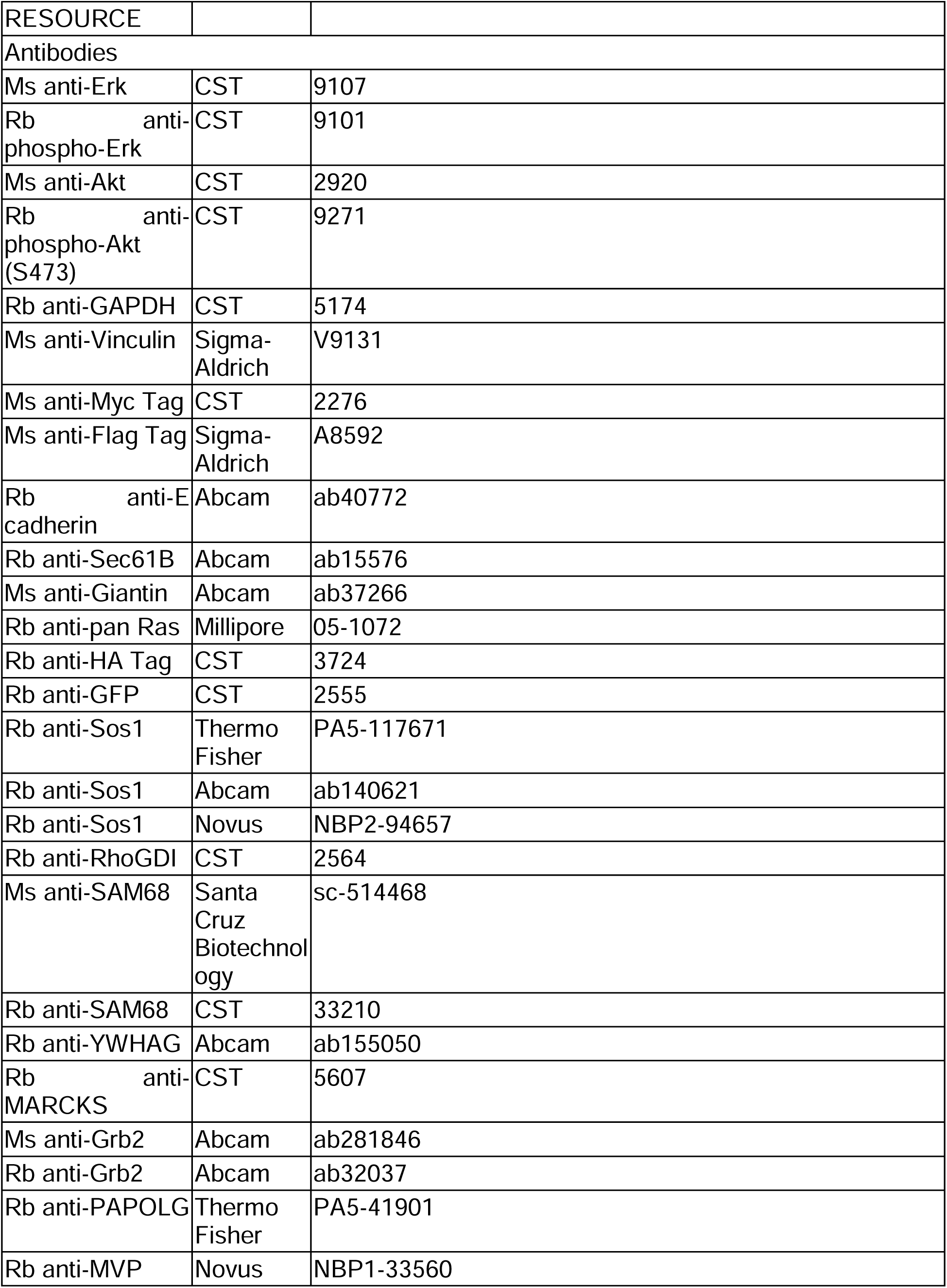
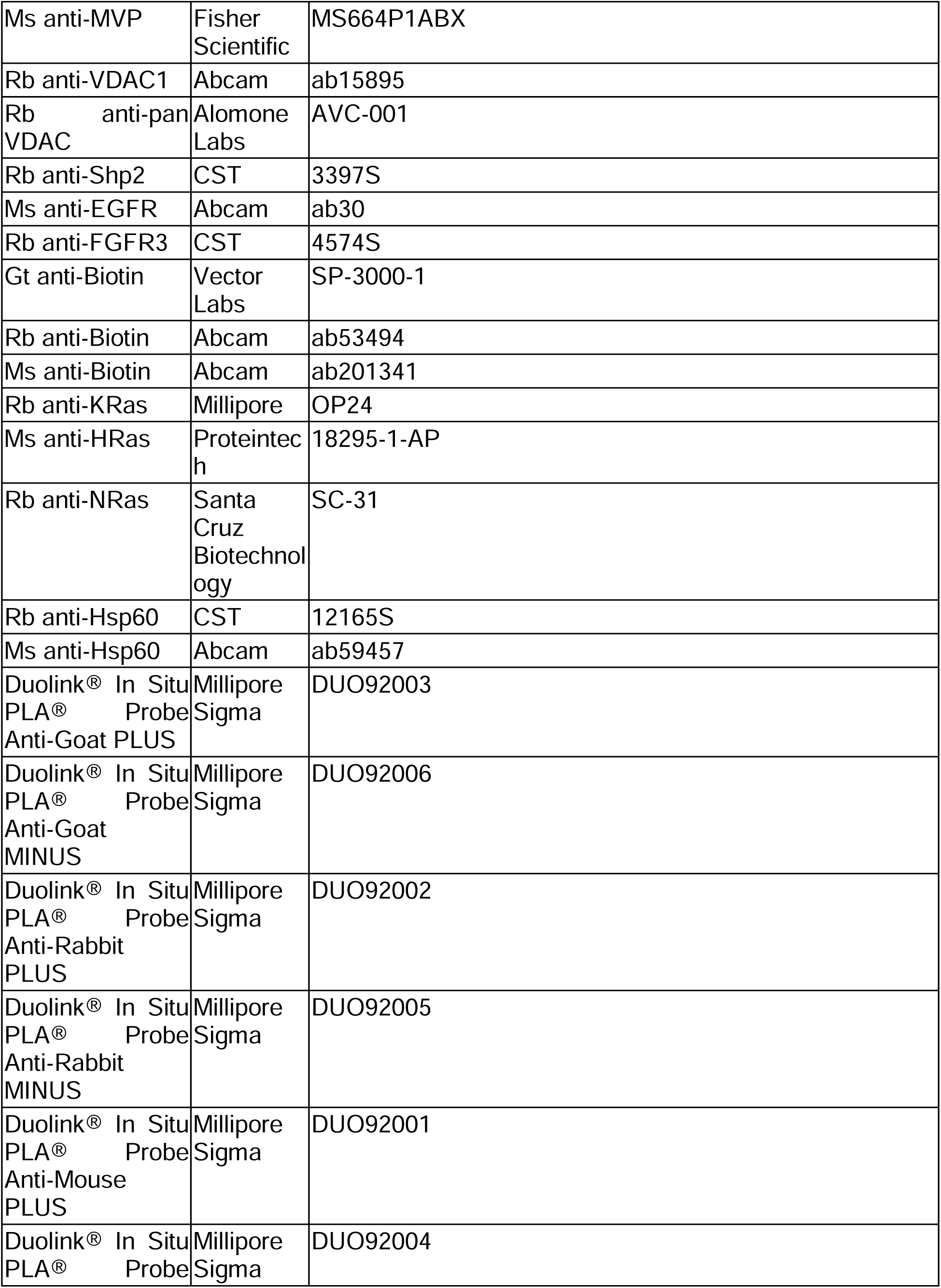
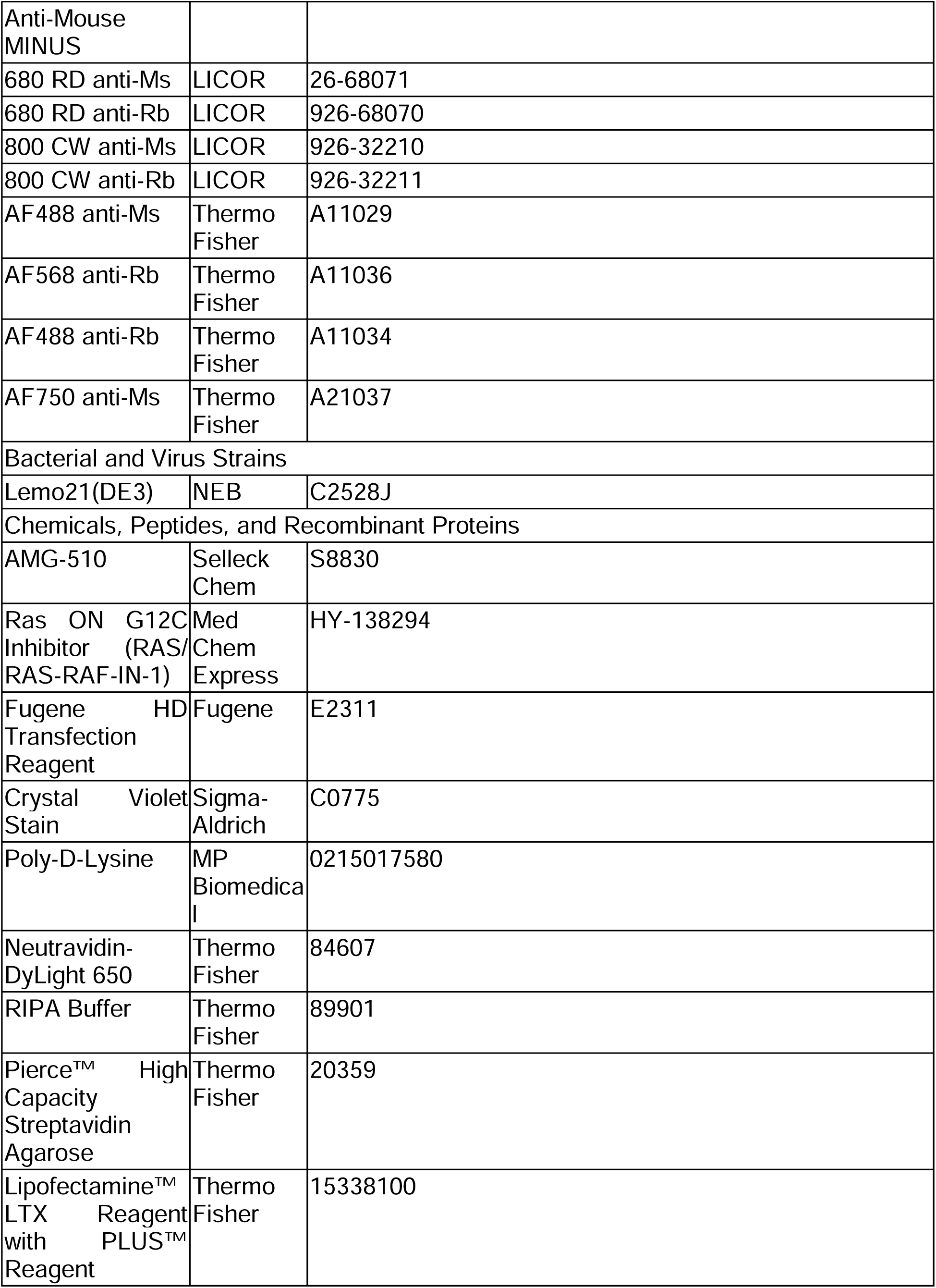
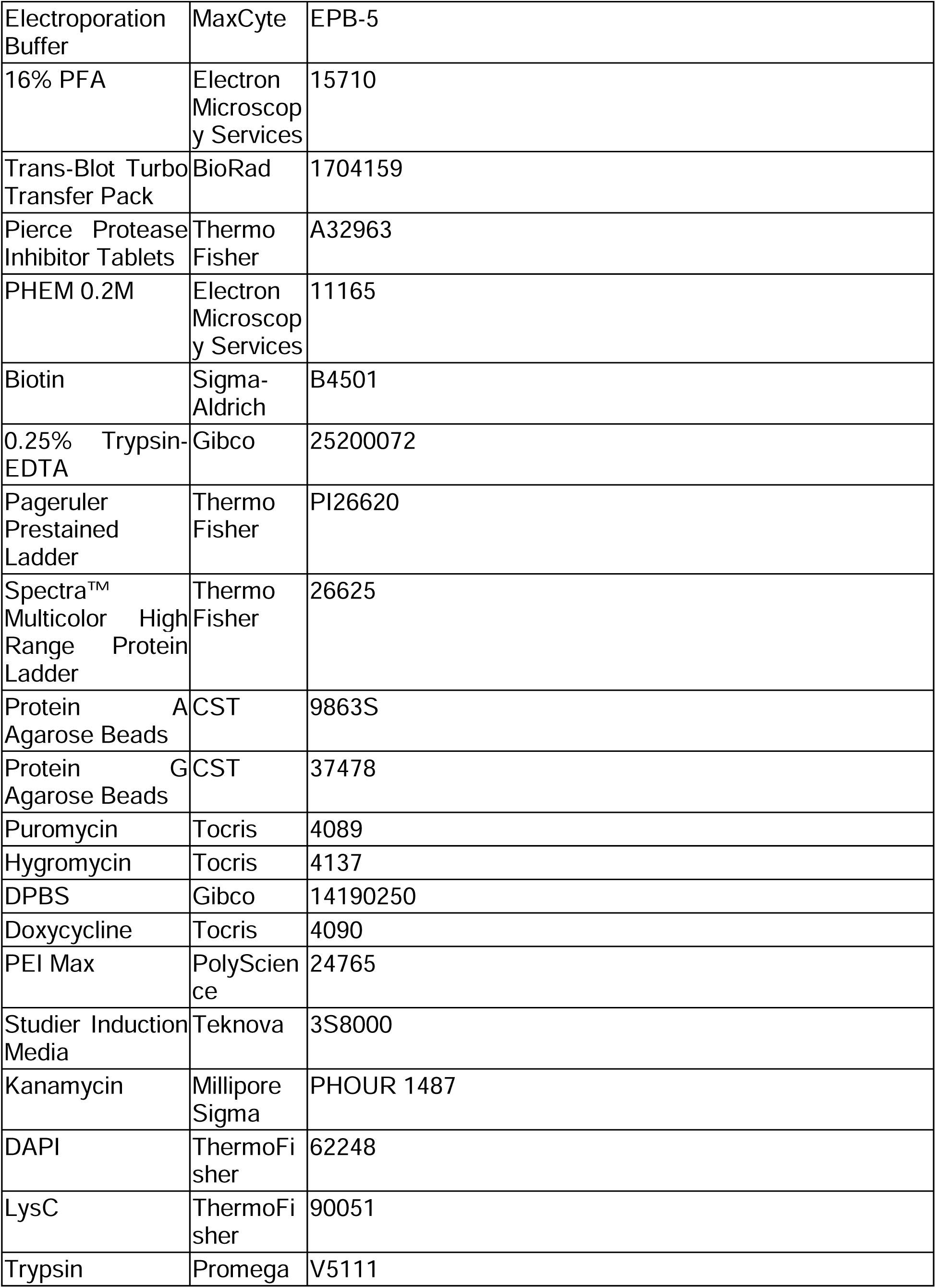
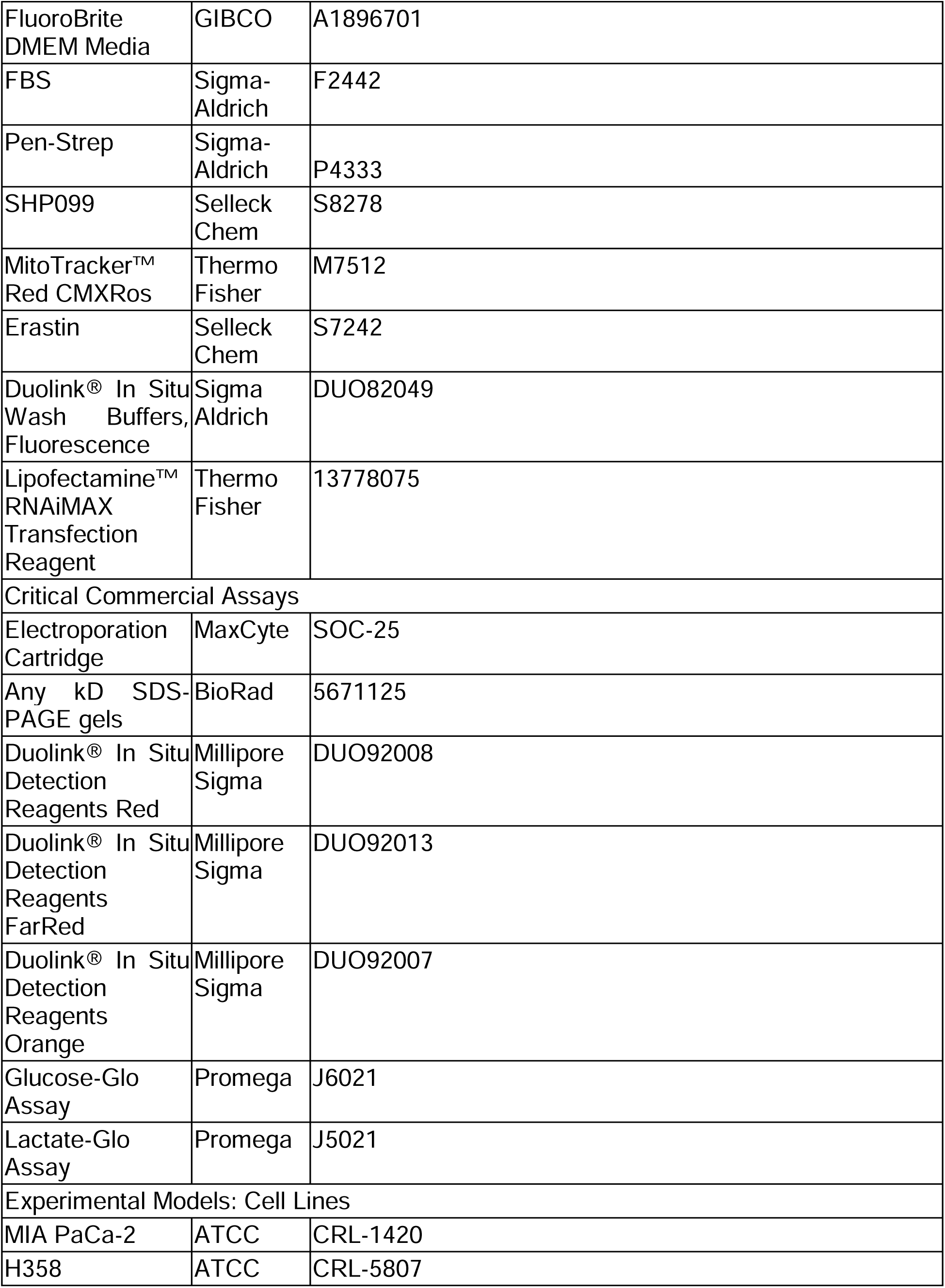
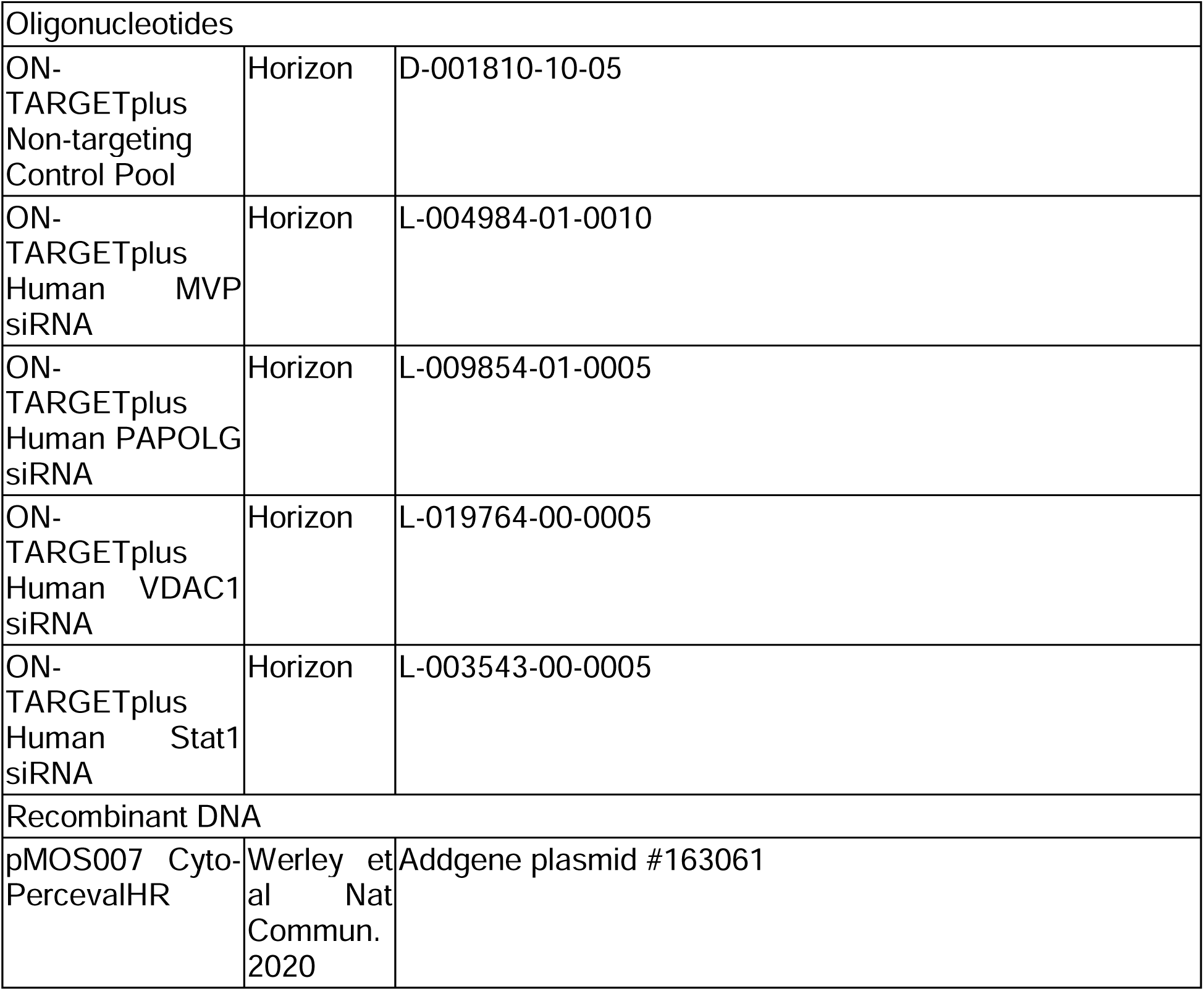

### RESOURCE AVAILABILITY

#### Lead contact

Further information and requests for resources and reagents should be directed to and will be fulfilled by the lead contact, Jason Zhang.

#### Materials Availability

Plasmids generated in this study can be shared upon request.

#### Data and code availability

The data that support the findings of this study are available from the corresponding author upon reasonable request. All accession codes have been provided for the paper. Source data are provided with this paper including mass spectrometry analysis. No additional code was generated for this study.

### EXPERIMENTAL MODEL AND SUBJECT DETAILS

#### Cell culture and transfection

MIA PaCa-2 and H358 cells were cultured in Dulbecco’s modified Eagle medium (DMEM) containing 1 g L^-1^ glucose and supplemented with 10% (v/v) fetal bovine serum (FBS) and 1% (v/v) penicillin–streptomycin (Pen-Strep). All cells were grown in a humidified incubator at 5% CO_2_ and at 37°C.

Before transfection, all cells were plated onto sterile poly-D-lysine coated plates or dishes and grown to 50%–70% confluence. siRNA transfection used Lipofectamine RNAiMAX. Besides siRNA transfection, the remaining transfections used Fugene HD and grown for an additional 16-24 hour before subsequent experiments. All cells underwent serum starvation for at least 16 hours before any treatment, unless indicated.

#### Generation of stable cell lines

Generating localized RasS17N in H358 and MIA-PaCa2 cells involved transfection of a pcDNA5 recombinase plasmid and an attb-containing plasmid at a 1:1 ratio that encodes for transgene and puromycin resistance gene. A negative control plate was also grown that was not transfected with the attb-containing plasmid. After cells were 10-25% confluent (usually 48 hours later), 100 ng mL^-1^ of hygromycin was added until 95% cell death in the negative control plate. Cells were recovered without hygromycin until cells are 80% confluent. Afterwards, recombined stable cell lines were maintained in 1 µg mL^-1^ puromycin.

### METHOD DETAILS

#### Plasmid construction

All plasmids constructed here are using the pcDNA 3.1 backbone (unless otherwise indicated) and were produced by GenScript.

#### Cell counting to measure cell proliferation

H358 and MIA PaCa-2 cells were seeded in 6-wells plates at 10,000 cells/well. After 1 day of plating, cells were treated with indicated drugs. Cell numbers were quantified using a hemacytometer each day for 7 days.

#### Colony formation assay

H358 and MIA PaCa-2 cells were seeded in 24-well plates at 100 cells/well. After 1 day of plating, cells were treated with indicated drugs. After 1-2 weeks to allow cell growth, cells were washed once with PBS, fixed with 4% paraformaldehyde (PFA) in PBS for 10 min, stained with 2.5 mg/mL crystal violet stain dissolved in 20% methanol for 10 min, and then washed 6x with PBS. Images were captured using the ZOE Fluorescent Cell Imager (BioRad).

#### Immunostaining and flow cytometry

H358 and MIA PaCa-2 cells were seeded onto 24-well glass-bottom plates. After transfection and drug addition, cells were fixed with 4% PFA in 2x PHEM buffer (60 mM PIPES, 50 mM HEPES, 20 mM EGTA, 4 mM MgCl_2_, 0.25 M sucrose, pH 7.3) for 10 min, permeabilized with 100% methanol for 10 min, washed with PBS 3x, blocked in 1% BSA in PBS for 30 min, incubated with primary antibody overnight at 4°C, washed with PBS 3x, incubated with DAPI and secondary antibodies for 1 hour at room temperature and aluminum foil cover. Cells were then washed with PBS 3x and mounted for epifluorescence imaging. All images were analyzed in ImageJ.

#### Proximity Ligation Assay

H358 and MIA PaCa-2 cells were seeded onto 24-well glass-bottom plates. After transfection and drug addition, cells were fixed with 4% PFA in 2x PHEM buffer for 10 min, permeabilized with 100% methanol for 10 min, washed with PBS 3x, blocked in 1% BSA in PBS for 30 min, incubated with primary antibody overnight at 4°C, washed with Wash Buffer A 2x, incubated with DAPI and secondary antibody with PLUS or MINUS DNA for 1 hour at 37°C, washed with Wash Buffer A 2x, incubated in ligation buffer for 30 min at 37°C, washed with Wash Buffer A 2x, incubated in amplification buffer for 100 min at 37°C, and finally washed with Wash Buffer B 3x. Cells were then mounted for epifluorescence imaging. All images were analyzed in ImageJ.

#### Immunoblotting and immunoprecipitation

Cells expressing indicated constructs and incubated with indicated drugs were plated, transfected, and labeled as described in figure legends. Cells were then transferred to ice and washed 2x with ice cold DPBS. Cells were then detached from the well by addition of 1x RIPA lysis buffer (50 mM Tris pH 8, 150 mM NaCl, 0.1% SDS, 0.5% sodium deoxycholate, 1% Triton X-100, 1x protease inhibitor cocktail, 1 mM PMSF, 1mM Na_3_VO_4_, 1% NP-40) and either scraping of cells or rotation on shaker for 30 min at 4°C. Cells were then collected and vortexed for at least 5 s every 10 min for 20 min at 4°C. Cells were then collected and clarified by centrifugation at 20,000 rpm for 10 minutes at 4°C. The supernatant was collected and underwent Pierce BCA assay to quantify total protein amounts.

For immunoblotting, whole cell lysate protein amounts were normalized across samples in the same gel, mixed with 4x loading buffer prior to loading, incubated at 95°C for 5 min and then 4°C for 5 min, and separated on Any kDa SDS-PAGE gels. Proteins separated on SDS-page gels were transferred to nitrocellulose membranes via the TransBlot system (BioRad). The blots were then blocked in 5% milk (w/v) in TBST (Tris-buffered saline, 0.1% Tween 20) for 1 hour at room temperature. Blots were washed with TBST 3x then incubated with indicated primary antibodies in 1% BSA (w/v) in TBST overnight at 4°C. Blots were then washed with TBST 3x and incubated with LICOR dye-conjugated secondary antibodies (LICOR 680/800 or streptavidin-LICOR 800) in 1% BSA (w/v) in TBST for 1 hour at room temperature. The blots were washed with TBST 3x and imaged on an Odyssey IR imager (LICOR). Quantitation of Western blots was performed using ImageJ on raw images.

For immunoprecipitation, agarose beads were either preloaded with streptavidin (high capacity streptavidin beads) or loaded by 3x lysis buffer washes and then addition of 1mg ml^-1^ indicated antibodies at 4°C on orbital shaker for 3 hour. Beads were then washed 2x in lysis buffer. Whole cell lysate protein amounts were normalized across samples and protein samples were added to beads (at least 100µg per sample) either at room temperature for 1 hour for streptavidin beads or at 4°C on orbital shaker overnight. Beads were then washed 2x in lysis buffer and 1x in TBS and then mixed with 4x loading buffer sometimes containing 2mM biotin and 20mM DTT^40^ for streptavidin pulldowns. The remaining portion of the protocol is the same as immunoblotting.

#### Mass spectrometry analysis

Cells expressing indicated constructs and incubated with indicated drugs were plated, transfected, and labeled as described in figure legends. Cells were then transferred to ice and washed 2x with ice cold DPBS, detached from the well by addition of 1x RIPA lysis buffer (50 mM Tris pH 8, 150 mM NaCl, 0.1% SDS, 0.5% sodium deoxycholate, 1% Triton X-100, 1x protease inhibitor cocktail, 1 mM PMSF, 1mM Na_3_VO_4_, 1% NP-40) and scraping of cells, collected and vortexed for at least 5 s every 10 min for 20 min at 4°C, and collected and clarified by centrifugation at 20,000g for 10 minutes at 4°C. The supernatant was collected and underwent Pierce BCA assay to quantify total protein amounts.

50µL of high capacity streptavidin agarose beads were washed 2x in lysis buffer. Whole cell lysate protein amounts were normalized across samples and protein samples were added to beads (at least 100µg per sample) at room temperature for 1 hour. Beads were then washed 2x with lysis buffer, 1x with 1M KCl, 1x with 0.1M Na_2_CO_3_, 2x 2M urea, and 2x with TBS. Beads were re-suspended in 50µL of denaturing buffer (6M guanidinium chloride, 50mM Tris containing 5mM TCEP and 10mM CAM with TCEP and CAM added fresh every time), inverted a few times, and heated to 95°C for 5 min. The bead slurry was diluted with 50µL of 100mM TEAB and 0.8µg of LysC was added per sample with the pH adjusted to 8-9 using 1M NaOH. This mixture was agitated on a thermomixer at 37°C for 2 hour at 1400 rpm. Afterwards, samples were diluted 2x with 100µL of 100mM TEAB with 0.8µg of sequencing grade trypsin per sample with the pH adjusted to 8-9 using 1M NaOH. This mixture was agitated on a thermomixer at 37°C for 12-14 hour at 800 rpm. After overnight trypsinization, samples were diluted 2x with 200µL of Buffer A (5% acetonitrile with 0.1% TFA) containing 1% formic acid. These samples were inverted a few times and pH adjusted to 2-3 using 100% formic acid. StageTips for peptide desalting were prepared by extracting out plugs from C18 matrices, shoved down a 200µL tip, and pressed with a plunger for flatness. Using these StageTips, 50µL of Buffer B (80% acetonitrile with 0.1% TFA) was passed through at 4000g for 1 min followed by 5µ0L of Buffer A for 4000g for 1 min. The supernatant of the samples was added to StageTips and spun down at 4000g for 5 min. Then, 50µL of Buffer A was added and spun down at 4000g for 2.5 min. 50µL of Buffer B was added to stage tips and a syringe pump was applied to elute samples.

Peptide samples were separated on an EASY-nLC 1200 System (Thermo Fisher Scientific) using 20 cm long fused silica capillary columns (100 µm ID, laser pulled in-house with Sutter P-2000, Novato CA) packed with 3 μm 120 Å reversed phase C18 beads (Dr. Maisch, Ammerbuch, DE). The LC gradient was 90 min long with 5−35% B at 300 nL/min. LC solvent A was 0.1% (v/v) aqueous acetic acid and LC solvent B was 20% 0.1% (v/v) acetic acid, 80% acetonitrile. MS data was collected with a Thermo Fisher Scientific Orbitrap Fusion Lumos using a data-dependent data acquisition method with a Orbitrap MS1 survey scan (R=60K) and as many Orbitrap HCD MS2 scans (R=30K) possible within the 2 second cycle time.

#### Computation of MS raw files

Data .raw files were analyzed by MaxQuant/Andromeda version 1.5.2.8 using protein, peptide and site FDRs of 0.01 and a score minimum of 40 for modified peptides, 0 for unmodified peptides; delta score minimum of 17 for modified peptides, 0 for unmodified peptides. MS/MS spectra were searched against the UniProt human database (updated July 22nd, 2015). MaxQuant search parameters: Variable modifications included Oxidation (M) and Phospho (S/T/Y). Carbamidomethyl (C) was a fixed modification. Max. missed cleavages was 2, enzyme was Trypsin/P and max. charge was 7. The MaxQuant “match between runs” feature was enabled. The initial search tolerance for FTMS scans was 20 ppm and 0.5 Da for ITMS MS/MS scans.

#### MaxQuant output data processing

MaxQuant output files were processed, statistically analyzed and clustered using the Perseus software package v1.5.6.0. Human gene ontology (GO) terms (GOBP, GOCC and GOMF) were loaded from the ‘mainAnnot.homo_sapiens.txt’ file downloaded on 02.03.2020. Expression columns (protein and phosphopeptide intensities) were log2 transformed and normalized by subtracting the median log2 expression value from each expression value of the corresponding data column. Potential contaminants, reverse hits and proteins only identified by site (biotinylation) were removed. Reproducibility between LC-MS/MS experiments was analyzed by column correlation (Pearson’s r) and replicates with a variation of r > 0.25 compared to the mean r-values of all replicates of the same experiment (cell line or knockdown experiment) were considered outliers and excluded from the analyses. Data imputation was performed in Perseus using a modeled distribution of MS intensity values downshifted by 1.8 and having a width of 0.2. Hits were further filtered using gene ontology analysis (signaling pathways) via Panther database.

#### Glucose consumption and lactate release assays

H358 and MIA PaCa-2 cells were seeded on 24-well plates at 10,000 cells/well. Supernatant fractions were collected after 1 day of cell settling and 72 hours after AMG-510 addition. Metabolite concentrations were quantified using the Lactate-Glo Assay and Glucose-Glo Assay, according to manufacturer’s instructions. Glucose consumption and lactate production were normalized to total protein content, which was measured by BCA protein assay.

#### Time-lapse epifluorescence imaging

Cells were washed twice with FluoroBrite DMEM imaging media and subsequently imaged in the same media in the dark at room temperature. Drugs were added for the indicated time durations. Epifluorescence imaging was performed on a Yokogawa CSU-X1 microscope with either a Lumencor Celesta light engine with 7 laser lines (408, 445, 473, 518, 545, 635, 750 nm) or a Nikon LUN-F XL laser launch with 4 solid state lasers (405, 488, 561, 640 nm), 40x/0.95 NA objective and a Hamamatsu ORCA-Fusion scientific CMOS camera, both controlled by NIS Elements 5.30 software (Nikon). The following excitation/emission filter combinations (center/bandwidth in nm) were used: CFP: EX445 EM483/32, CFP/YFP FRET: EX445 EM542/27, YFP: EX473 EM544/24, GFP: EX473 EM525/36, RFP: EX545 EM605/52, Far Red (e.g. AlexaFluor 647): EX635 EM705/72, 450: EX445 EM525/36, 500: EX488 EM525/36. Exposure times were 100 ms for acceptor direct channel and 500ms for all other channels, with no EM gain set and no ND filter added. Cells that were too bright (acceptor channel intensity is 3 standard deviations above mean intensity across experiments) or with significant photobleaching prior to drug addition were excluded from analysis. All epifluorescence experiments were subsequently analyzed using Image J. Brightfield images were acquired on the ZOE Fluorescent Cell Imager (BioRad).

### QUANTIFICATION AND STATISTICAL ANALYSIS

#### FRET biosensor analysis

Raw fluorescence images were corrected by subtracting the background fluorescence intensity of a cell-free region from the emission intensities of biosensor-expressing cells. Cyan/yellow FRET ratios were then calculated at each time point (*R)*. For some curves, the resulting time courses were normalized by dividing the FRET ratio at each time point by the basal ratio value at time zero (*R*/*R*_0_), which was defined as the FRET ratio at the time point immediately preceding drug addition (*R*_0_). Graphs were plotted using GraphPad Prism 8 (GraphPad).

#### Co-localization analysis

For co-localization analysis, cell images were individually thresholded and underwent Coloc 2 analysis on ImageJ.

#### Quantification of PLA puncta

For analysis of puncta number, cell images were individually thresholded and underwent particle analysis with circularity and size cutoffs in ImageJ.

#### Statistics and reproducibility

No statistical methods were used to predetermine the sample size. No sample was excluded from data analysis, and no blinding was used. All data were assessed for normality. For normally distributed data, pairwise comparisons were performed using unpaired two-tailed Student’s t tests, with Welch’s correction for unequal variances used as indicated. Comparisons between three or more groups were performed using ordinary one-way or two-way analysis of variance (ANOVA) as indicated. For data that were not normally distributed, pairwise comparisons were performed using the Mann-Whitney U test, and comparisons between multiple groups were performed using the Kruskal-Wallis test. All data shown are reported as mean and error bars in figures represent ± SEM. All data was done at least in biological triplicates unless otherwise stated. All data were analyzed and plotted using GraphPad Prism 8 including non-linear regression fitting. Coefficient of variation (CV) values are calculated as standard deviation divided by mean.

## Acknowledgements

We acknowledge funding from HHMI (J.Z.Z. and D.B.), Helen Hay Whitney Foundation (J.Z.Z.), the Audacious Project at the Institute for Protein Design (J.Z.Z, D.B.), NIH grants (R01GM129090 (S-E.O.), R01GM145011 (D.J.M.), and R01GM086858 (D.J.M.)). K. Shokat for fruitful discussion of the Ras results.

## Author contributions

J.Z.Z. conceived and supervised the project. J.Z.Z. and D.J.M. designed the experiments. J.Z.Z. and W.H.N. performed all experiments. S.E.O. ran samples through mass spectrometry. J.Z.Z. and D.J.M. wrote the original draft. All authors reviewed and commented on the manuscript.

## Inclusion and diversity

One or more of the authors of this paper self-identifies as a member of the LGBTQIA+ community. While citing references scientifically relevant for this work, we also actively worked to promote gender balance in our reference list.

## Figure Legends

**Figure S1:**
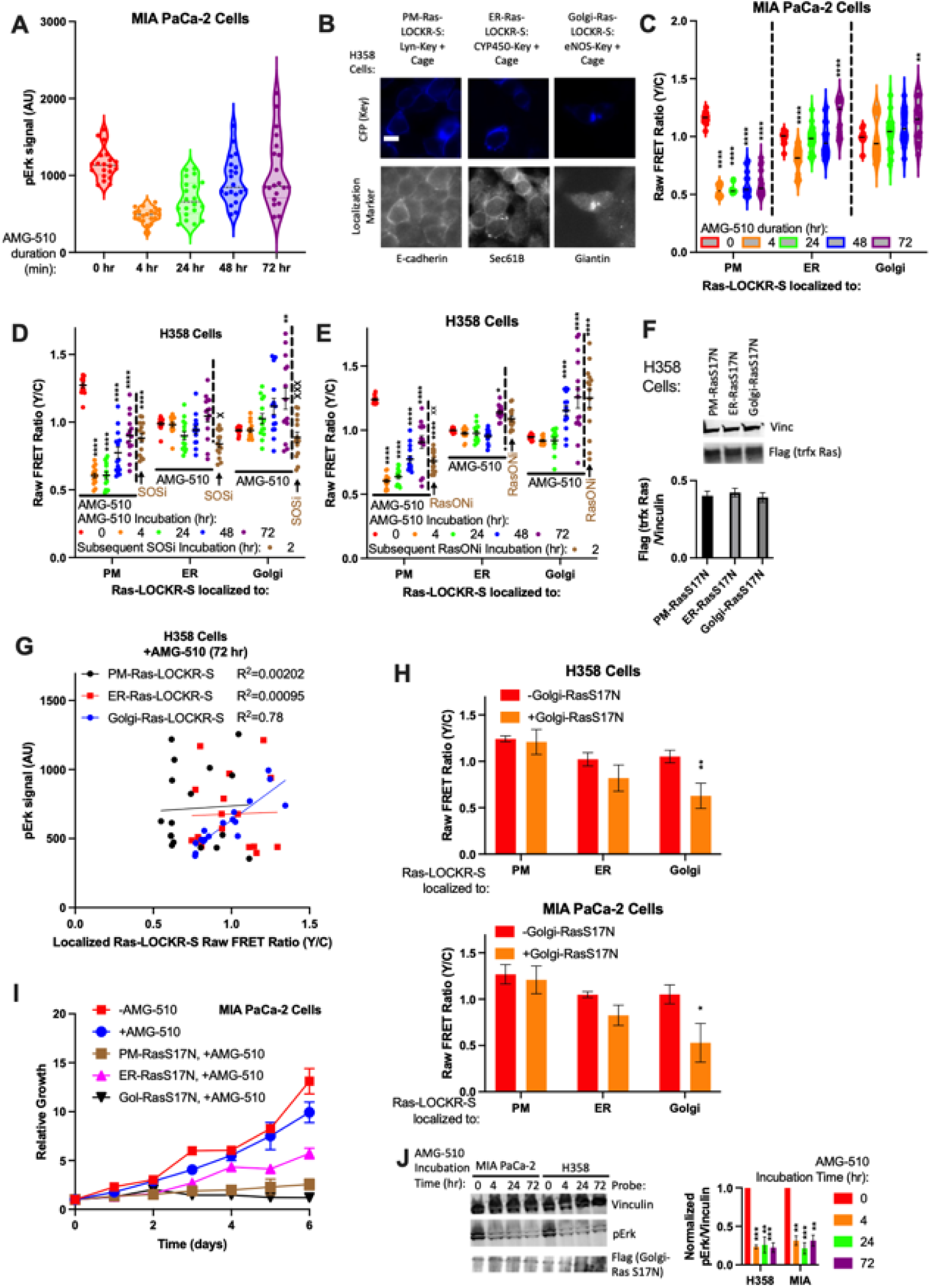
RasG12C-GDP inhibitor treatment leads to golgi-localized Ras activity that drives oncogenic signaling and growth, Related to Figure 1. (A) Violin plots with SEM values on bottom of MIA PaCa-2 cells treated with 100nM AMG-510 over time and immunostained for pErk. Each dot represents the average fluorescence intensity (from pErk immunostaining) of single cell (n=20 cells per condition). Statistics: Ordinary one-way ANOVA, comparison to 0 hour AMG-510 time point. CV at 0 hour=17%, 4 hour=20%, 24 hour=35%, 48 hour=33%, 72 hour=44%. (B) H358 cells transfected with Ras-LOCKR-S localized to different subcellular regions were immunostained for a protein known to localize to same region. Scale bar = 10µm. (C) Raw FRET ratios of localized Ras-LOCKR-S in MIA PaCa-2 cells treated over time with 100nM AMG-510 (n=17 cells per condition). PM: CV at 0 hour=4.1%, 4 hour=9.4%, 24 hour=6.4%, 48 hour=18%, 72 hour=19%. ER: CV at 0 hour=5.2%, 4 hour=14%, 24 hour=12%, 48 hour=15%, 72 hour=13%. Golgi: CV at 0 hour=7.4%, 4 hour=18%, 24 hour=16%, 48 hour=14%, 72 hour=16% (D-E) Raw FRET ratios (Y/C) of localized Ras-LOCKR-S in H358 cells treated over time with 100nM AMG-510 (n=17 cells per condition). After 72 hour AMG-510 incubation, either 1μM Sos inhibitor BI-3406 (D) or 100nM RasG12C-GTP (RasON) inhibitor (RasONi) RMC-6291 (E) was added for 2 hours. Statistics: Ordinary two-way ANOVA. * is comparison to 0 hour AMG-510 time point and x is comparison to 72 hour AMG-510 time point. (F) Left: Representative immunoblot of H358 cells expressing RasS17N localized to different subcellular regions (n=3 experiments). Right: Densitometry analysis with bar representing average and error bars representing SEM. (G) Scatterplot comparing localized Ras-LOCKR-S raw FRET ratio to pErk immunostaining fluorescence in H358 cells treated with 100nM AMG-510 for 72 hours. Each dot represents a single cell where both measurements were done (n=17 cells per condition). (H) Average raw FRET ratios of H358 or MIA PaCa-2 cells transfected localized Ras-LOCKR-S with or without golgi-localized RasS17N (n=15 cells per condition). Bars represent average and error bars represent SEM. Statistics: Ordinary two-way ANOVA, comparison to 0 hour AMG-510 time point. (I) Cell growth curves of MIA PaCa-2 cells expressing localized RasS17N and/or treatment of 100nM AMG-510 (n=3 experiments per condition). Line represents average from all 3 experiments. (J) Left: Representative immunoblot of MIA PaCa-2 or H358 cells expressing localized RasS17N and treated with 100nM AMG-510 for various time durations (n=4 experiments). Right: Densitometry analysis with bar representing average and error bars representing SEM. Data is normalized to no incubation (0 hour time point). Statistics: Ordinary two-way ANOVA, comparison to 0 hour AMG-510 time point.

**Figure S2:**
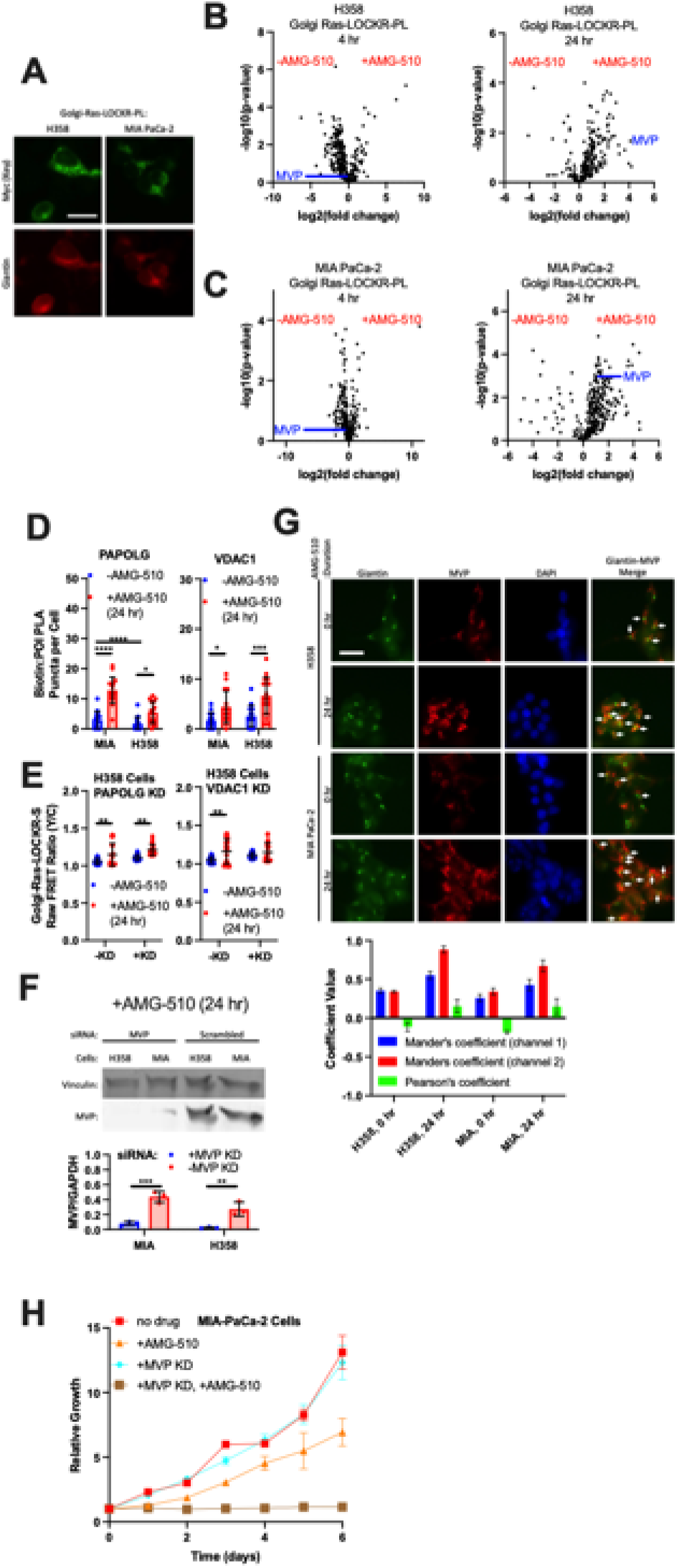
Identifying MVP as a mediator of oncogenic signaling and cell growth during AMG-510 treatment, Related to Figure 2. (A) Representative epifluorescence images of H358 or MIA PaCa-2 cells transfected with golgi-localized Ras-LOCKR-PL and immunostained for a golgi marker, giantin. Scale bar = 10µm. (B-C) Volcano plots of mass spectrometry results of H358 (B) or MIA PaCa-2 (C) cells expressing localized Ras-LOCKR-PL and stimulated for 24 hours with 500µM biotin and either with or without 100nM AMG-510. Plotted differences are comparing with AMG-510 (right) or without AMG-510 (left). MVP is highlighted in blue. (D) MIA PaCa-2 or H358 cells were transfected with golgi-localized Ras-LOCKR-PL, incubated with 500µM biotin and either with DMSO or 100nM AMG-510, and underwent PLA immunostaining (n=16 cells). Shown is average amount of biotin:PAPOLG or biotin:VDAC1 PLA puncta (anti-biotin, anti-PAPOLG, anti-VDAC1) per cell. Each dot represents a single cell, bars represent average, and error bars represent SEM. Statistics: Student’s t-test. (E) Comparison of raw FRET ratios of localized Ras-LOCKR-S between no KD versus PAPOLG KD or VDAC1 KD in H358 cells treated with 100nM AMG-510 over indicated times (n=17 cells per condition). Statistics: Student’s t-test. (F) Top: Representative immunoblot of MIA PaCa-2 or H358 cells transfected with MVP or scrambled siRNA for 2 days and treated with 100nM AMG-510 for 1 day. Bottom: Quantification across conditions (n=3 experiments per condition). Statistics: Student’s t-test. (G) Representative epifluorescence images of H358 and MIA PaCa-2 cells treated without (0 hour) or with 100nM AMG-510 (24 hour) and immunostained for MVP and a localization marker for the golgi (Giantin). Colocalization analysis is shown as well (n=3 experiments per condition). (H) Cell growth curves of MIA PaCa-2 cells with MVP KD and/or treatment of 100nM AMG-510 (n=3 experiments per condition). Line represents average from all 3 experiments.

**Figure S3:**
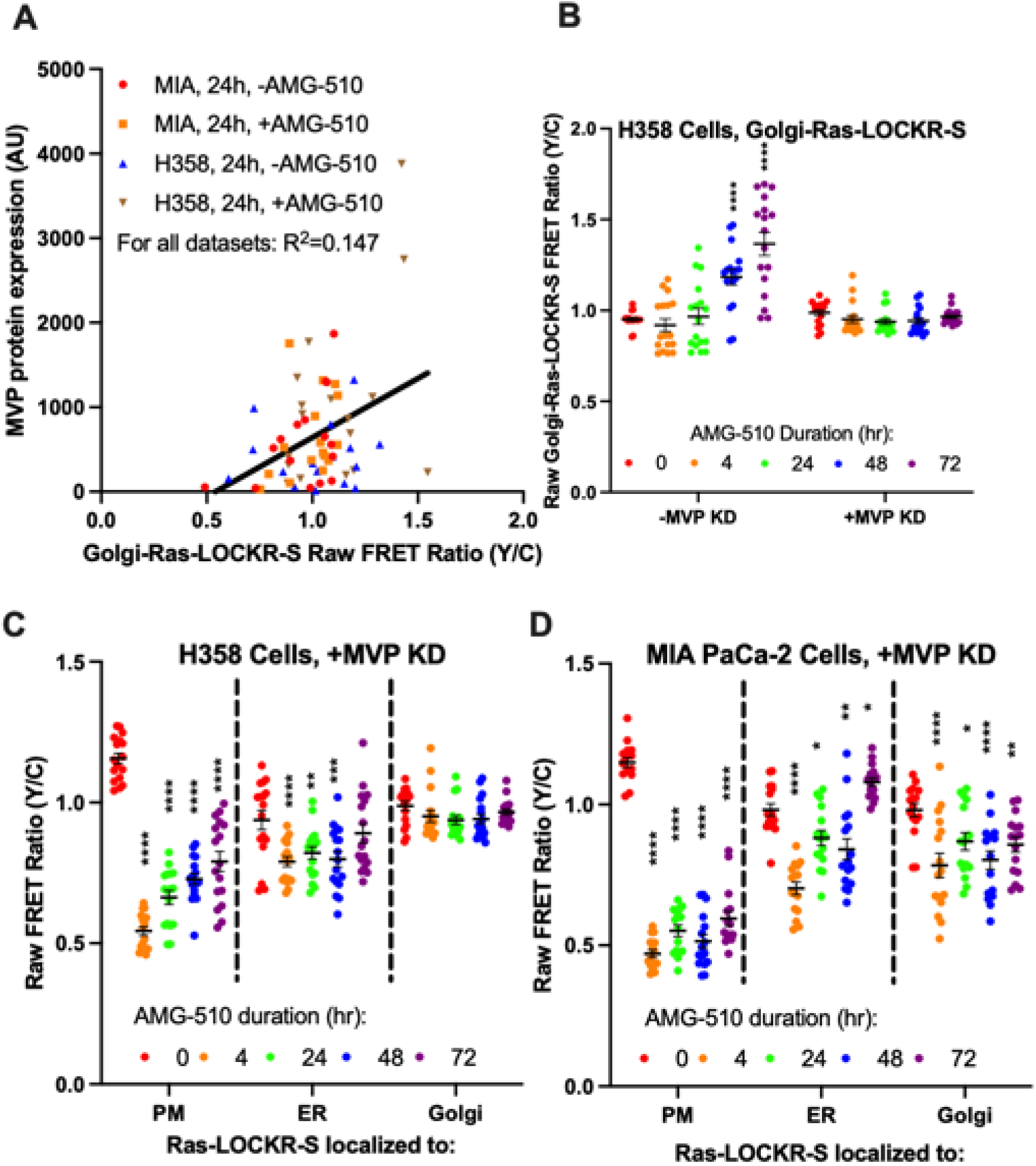
MVP at the golgi-Ras signalosome typifies the Amg-510-promoted signaling adaptive cell subpopulation, Related to Figure 3. (A) Scatterplot comparing MVP expression by immunostaining (y-axis) to golgi-Ras-LOCKR-S raw FRET ratios (x-axis). MIA PaCa-2 or H358 cells were transfected with golgi-Ras-LOCKR-S, treated either with DMSO or 100nM AMG-510, immunostained, and then imaged. Each dot represents a single cell where both measurements were done (n=16 cells per condition). (B) Raw FRET ratios of golgi-Ras-LOCKR-S in H358 cells transfect either with scrambled or MVP siRNA and then treated over time with 100nM AMG-510 (n=17 cells per condition). Statistics: Ordinary two-way ANOVA, comparison to 0 hour AMG-510 time point (C-D) Raw FRET ratios of subcellularly localized Ras-LOCKR-S in either H358 (C) or MIA PaCa-2 (D) cells with MVP KD and then treated over time with 100nM AMG-510 (n=17 cells per condition). Statistics: Ordinary two-way ANOVA, comparison to 0 hour AMG-510 time point.

**Figure S4:**
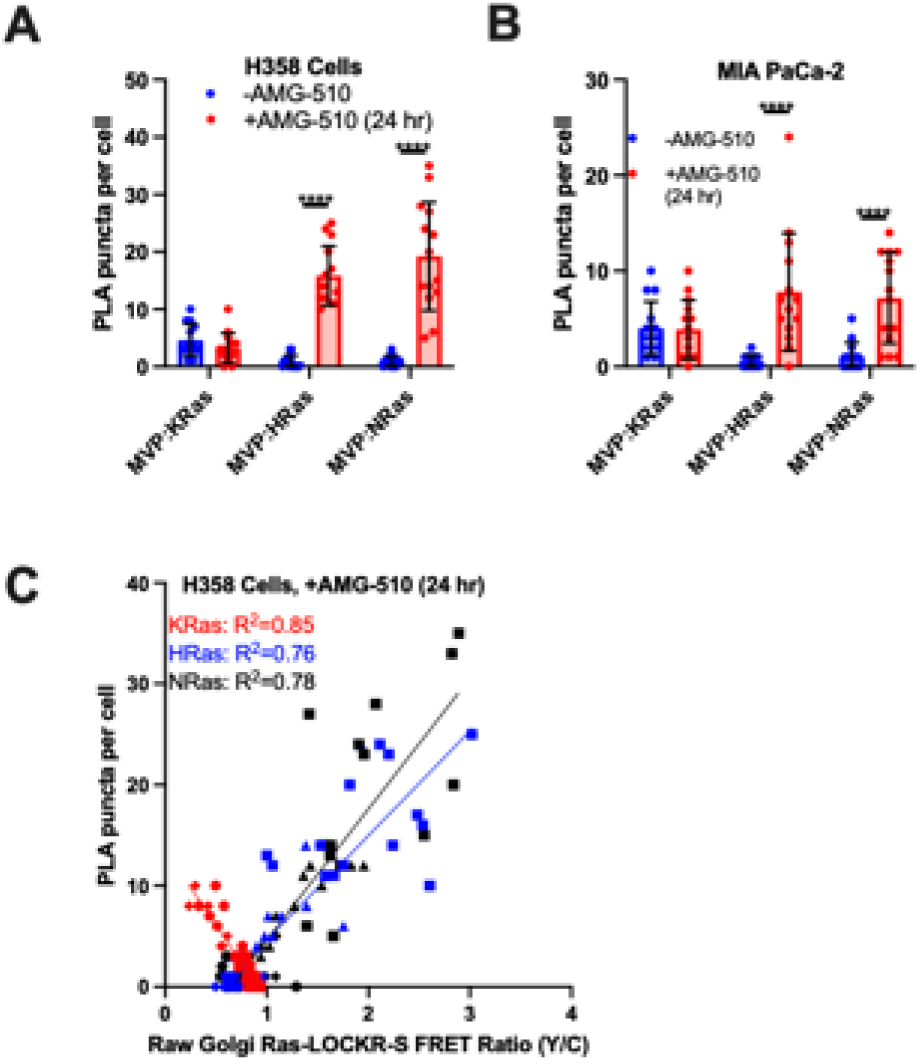
MVP interactions with Ras isoforms change during AMG-510 inhibition, Related to Figure 4. (A-C) H358 (A) or MIA PaCa-2 (B) cells transfected with golgi-Ras-LOCKR-S and treated with either DMSO or 100nM AMG-510 for 24 hours were immunostained and probed for MVP:Ras isoform interactions via PLA. Displayed is (A-B) quantification across conditions and (C) correlation between golgi-Ras-LOCKR-S raw FRET ratios and PLA puncta (anti-MVP, anti-KRas, anti-HRas, anti-NRas) per cell (n=14 cells per condition). Same dataset is used in **Fig. S4A-B**. Statistics: Student’s t-test.

**Figure S5:**
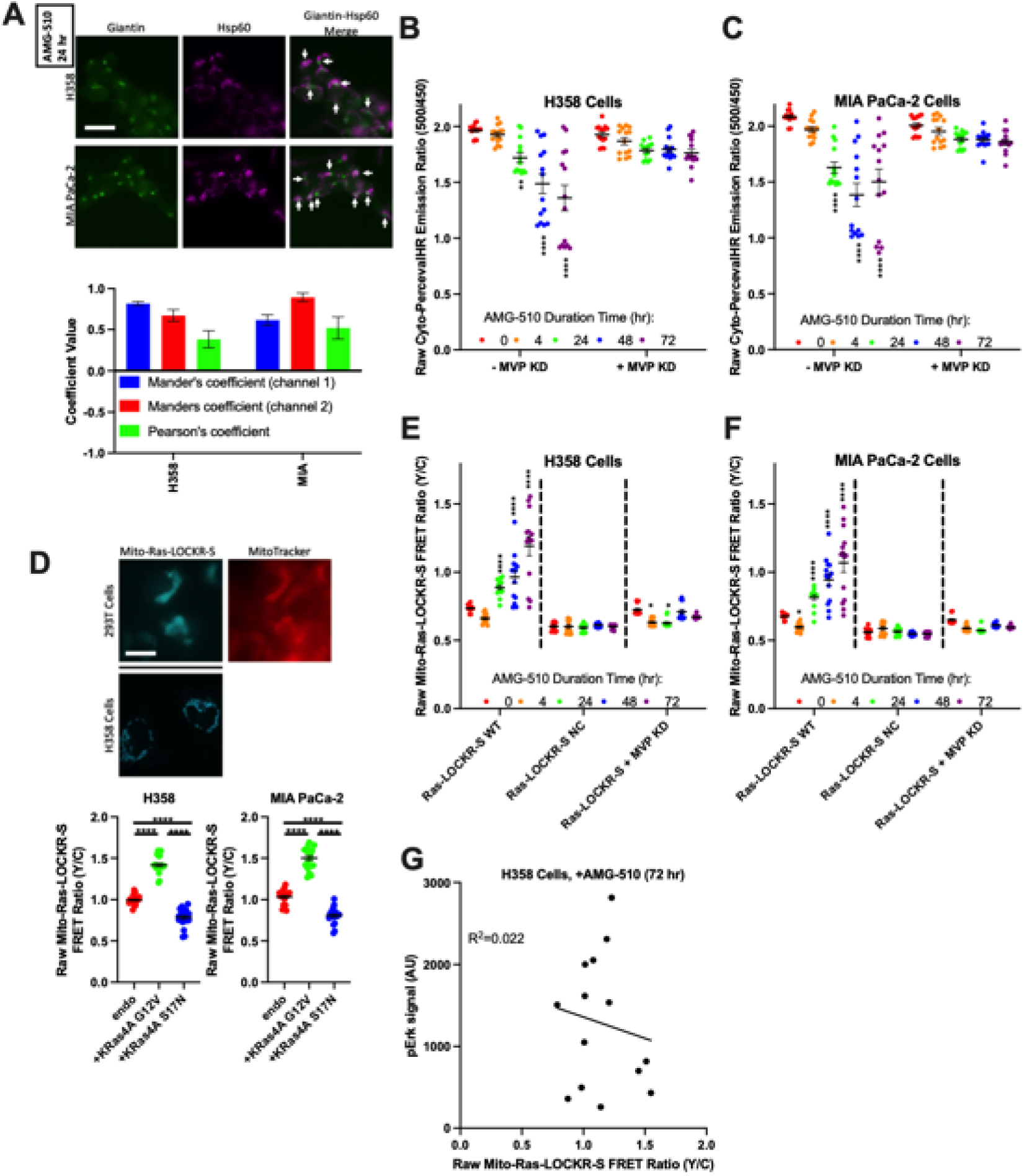
MVP drives metabolic changes during AMG-510 treatment, Related to Figure 5. (A) Top: Representative epifluorescence images of H358 and MIA PaCa-2 cells treated with 100nM AMG-510 for 24 hours and immunostained for golgi (Giantin) and mitochondria (Hsp60) localization markers. Bottom: Colocalization analysis (n=3 experiments per condition) (B-C) Raw emission ratios of H358 (B) or MIA PaCa-2 (C) cells transfected with cyto-PercevalHR and either scrambled or MVP siRNA and treated over time with 100nM AMG-510 (n=14 cells per condition). Statistics: Ordinary two-way ANOVA. (D) Top: Representative epifluorescence image of 293T or H358 cells transfected with mito-Ras-LOCKR-S and probed with MitoTracker Red. Bottom: Raw FRET ratios of H358 and MIA PaCa-2 cells transfected with mito-Ras-LOCKR-S and either transfected with nothing else, GFP-tagged HRas G12V, or GFP-tagged HRas S17N (n=20 cells per condition). (E-F) Raw FRET ratios of H358 (E) or MIA PaCa-2 (F) transfected with either mito-Ras-LOCKR-S WT, mito-Ras-LOCKR-S NC sensor, or mito-Ras-LOCKR-S WT and MVP siRNA, and treated over time with 100nM AMG-510 (n=14 cells per condition). Statistics: Ordinary two-way ANOVA. (G) Scatterplot comparing mito-Ras-LOCKR-S raw FRET ratio to pErk immunostaining fluorescence in H358 cells treated with 100nM AMG-510 for 72 hours. Each dot represents a single cell where both measurements were done (n=14 cells).

**Figure S6:**
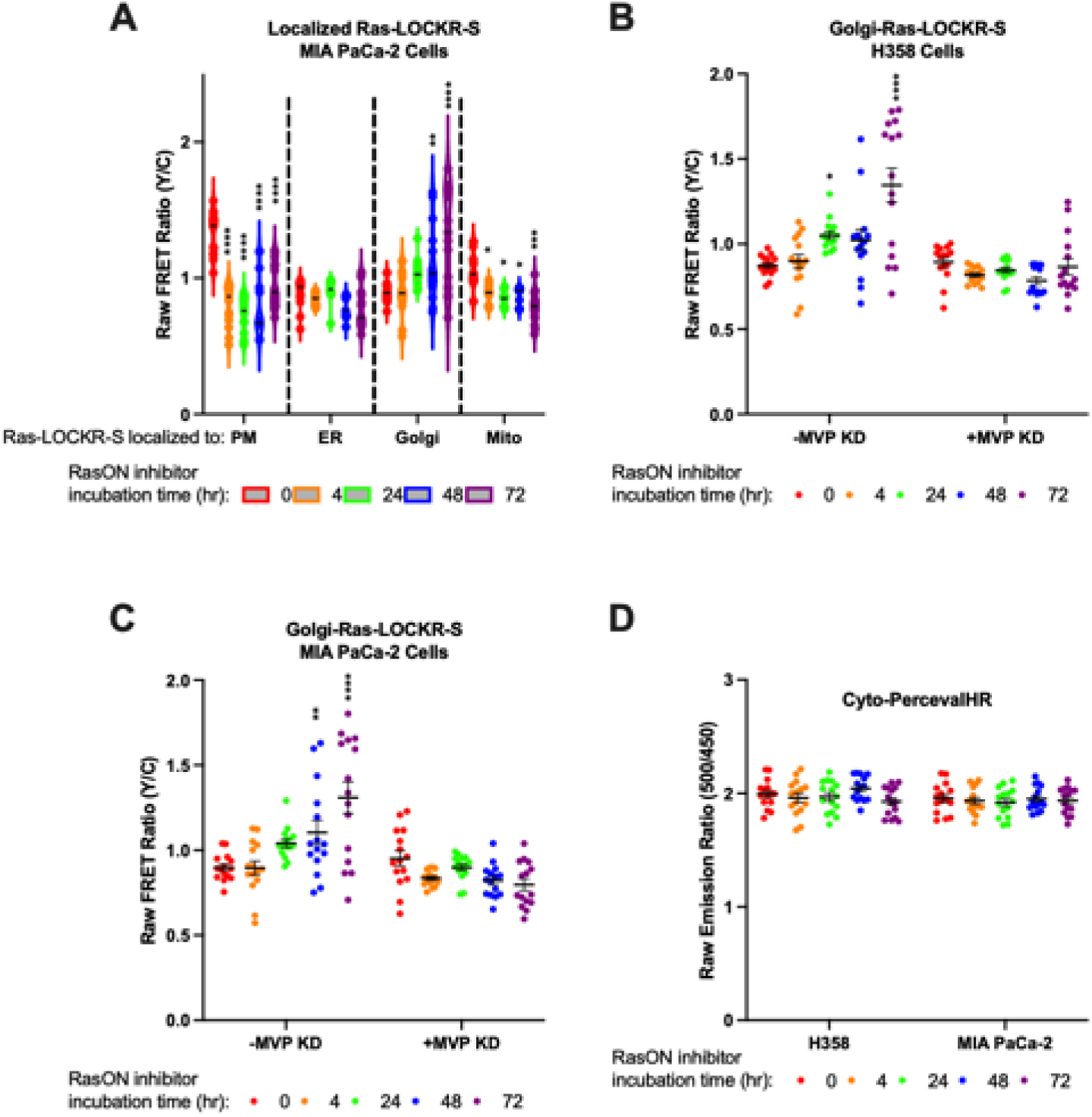
RasONi treatment induces compensatory oncogenic signaling that is dependent on MVP, Related to Figure 6. (A) Raw FRET ratios of MIA PaCa-2 cells transfected with subcellularly localized Ras-LOCKR-S and treated over time with 100nM RasONi (n=15 cells per condition). Statistics: Ordinary two-way ANOVA, comparison to 0 hour RasONi time point. PM: CV at 0 hour=12%, 4 hour=19%, 24 hour=18%, 48 hour=28%, 72 hour=18%. ER: CV at 0 hour=12%, 4 hour=5.2%, 24 hour=8.6%, 48 hour=10%, 72 hour=19%. Golgi: CV at 0 hour=8.6%, 4 hour=18%, 24 hour=8.8%, 48 hour=26%, 72 hour=28%. Mito: CV at 0 hour=13%, 4 hour=8.1%, 24 hour=6.8%, 48 hour=8.1%, 72 hour=15%. (B-C) Raw FRET ratios of H358 (B) or MIA PaCa-2 (C) cells transfected either with scrambled or MVP siRNA for 2 days, transfected with golgi-Ras-LOCKR-S, and treated over time with 100nM RasONi (n=15 cells per condition). Statistics: Ordinary two-way ANOVA, comparison to 0 hour RasONi time point. (D) Raw emission ratios of H358 and MIA PaCa-2 cells transfected with cyto-PercevalHR and treated over time with 100nM RasONi (n=15 cells per condition). Statistics: Ordinary two-way ANOVA, comparison to 0 hour RasONi time point.

## Table Legends

**Table S1: Differential golgi-Ras-LOCKR-PL labeling during AMG-510 treatment** H358 cells transfected with golgi-Ras-LOCKR-PL were incubated with 500µM biotin and with or without 100nM AMG-510 for either 4 hours or 24 hours. Afterwards, cells were lysed, underwent streptavidin pulldown, trypsin digested, and sent for MS analysis. Hits identified by MS were filtered based on selectively labeling (more than 2-fold change compared to no AMG-510), passed our statistical cutoff, and were related to signaling based on gene ontology analysis (see Methods for details).

